# Hearing loss in Alzheimer Disease is associated with altered serum lipidomic biomarker profiles

**DOI:** 10.1101/2020.08.15.252452

**Authors:** Daniel A. Llano, Lina K. Issa, Priya Devanarayan, Viswanath Devanarayan, Alzheimer’s Disease Neuroimaging Initiative (ADNI)

## Abstract

Recent data have found that aging-related hearing loss (ARHL) is associated with the development of Alzheimer Disease (AD). However, the nature of the relationship between these two disorders is not clear. There are multiple potential factors that link ARHL and AD, and previous investigators have speculated that shared metabolic dysregulation may underlie the propensity to develop both disorders. Here, we investigate the distribution of serum lipidomic biomarkers in AD subjects with or without hearing loss in a publicly available dataset. Serum levels of 349 known lipids from 16 lipid classes were measured in 185 AD patients. Using previously defined co-regulated sets of lipids, both age- and sex-adjusted, we found that lipid sets enriched in phosphatidylcholine and phosphatidylethanolamine showed a strong inverse association with hearing loss. Examination of biochemical classes confirmed these relationships and revealed that serum phosphatidylcholine levels were significantly lower in AD subjects with hearing loss. A similar relationship was not found in normal subjects. These data suggest that a synergistic relationship may exist between AD, hearing loss and metabolic biomarkers, such that in the context of a pathological state such as AD, alterations in serum metabolic profiles are associated with hearing loss. These data also point to a potential role for phosphatidylcholine, a molecule with antioxidant properties, in the underlying pathophysiology of ARHL in the context of AD, which has implications for our understanding and potential treatment of both disorders.

## Introduction

Aging-related hearing loss (ARHL) and Alzheimer Disease (AD) are common disabling disorders in the elderly. Over the age of 65, approximately 10% of individuals develop AD, while approximately 40% develop ARHL (Hebert, Weuve, Scherr, & Evans, 2013; Nash et al., 2011), Both disorders are rising in prevalence as the population ages, and an estimated 83 million individuals will be over the age of 65 in the U.S. by the year 2050 (Ortman, Velkoff, & Hogan, 2014). Recent data have revealed an association between AD and ARHL, such that the likelihood of developing cognitive impairment, and ultimately AD, is increased in individuals with ARHL (Ford et al., 2018; Golub et al., 2017; Lin, Ferrucci, et al., 2011; Lin, Metter, et al., 2011; Lin, Thorpe, Gordon-Salant, & Ferrucci, 2011; Lin et al., 2013; Panza, Solfrizzi, & Logroscino, 2015; Thomson, Auduong, Miller, & Gurgel, 2017). This relationship holds true even when adjusting for age, sex and multiple other potentially confounding variables such as comorbid illness. A causal association has not been identified, though multiple mechanisms by which hearing loss may lead to AD have been proposed (reviewed in (Nadhimi & Llano, 2020)).

ARHL and AD do share several potential biological substrates. Both are associated with metabolic stress and diminished mitochondrial function (Fujimoto & Yamasoba, 2014; Wang et al., 2014). ARHL is also associated with more traditional markers of AD such as increases in CSF tau and diminished hippocampal and entorhinal cortical volume (Xu et al., 2019). Recently, it has been suggested that AD may be associated with widespread dysregulation of lipid metabolism (Kao, Ho, Tu, Jou, & Tsai, 2020) and plasma lipid profiles have been shown to correlate with multiple AD-related biomarkers (Barupal et al., 2019). Lipid dysregulation may also play a role in the development of hearing loss (Campbell, Rybak, & Khardori, 1996). It is therefore possible that an underlying process of metabolic dysregulation, including altered lipid homeostasis, may account for the relationship between AD and ARHL.

Lipids are a major component of biological membranes and integral for neuronal function. The term lipids encompasses all fatty acids and their derivatives. Body lipids are derived from three sources: our diet, adipose tissue storage, and the liver’s synthetic capacity. Fats ingested in the diet enter the gastrointestinal tract, digested by pancreatic lipases in the small intestine and then imported across the intestinal mucosa. Lipids are then packaged along with cholesterol into chylomicrons which allow for nonpolar substances to move within the aqueous environment of our lymphatic and circulatory systems. These fats are then oxidized through β-oxidation for energy production or re-esterized for storage in adipose tissue. Alternatively, lipids in the small intestine can be distributed to the liver through portal circulation or to adipose tissue. Lipids derived from endogenous synthesis in the liver are packaged into very-low-density lipoproteins that are transported to tissue or stored in adipose tissue. Fat stores in adipose tissue are mobilized for energy production by the action of hormone-sensitive lipase as needed.

Given the potential roles for lipid dysregulation in the development of both AD and ARHL, and recently discovered associations between serum lipid profiles and AD pathological biomarkers (Barupal et al., 2019), we hypothesized that serum lipids may be disrupted in AD subjects with hearing loss. Therefore, in the current study, we examined the distribution of serum lipids in subjects with AD, with and without hearing loss, using a publicly-available dataset (Alzheimer Disease Neuroimaging Initiative, ADNI).

## Methods

### Database

The ADNI database (adni.loni.usc.edu) utilized in this research was launched in 2003 as a public-private partnership, led by Principal Investigator Michael W. Weiner, MD. The primary goal of ADNI has been to test whether serial MRI, PET, other biological markers, and clinical and neuropsychological assessments can be combined to measure the progression of mild cognitive impairment (MCI) and early AD. For up-to-date information, see www.adni-info.org. This study was conducted across multiple clinical sites and was approved by the Institutional Review Boards of all of the participating institutions. Informed written consent was obtained from all participants at each site. Data used for the analyses presented here were accessed on June 25, 2020.

### Lipid analysis

Details of lipid extraction and measurement as well as quality control measures have been previously described (Barupal et al., 2018). In brief, fasting serum samples were obtained from subjects during the baseline visit. Lipids were extracted using organic solvents. Serum extracts were then analyzed using liquid chromatography with mass spectrometry. After quality control measures, data were available from a total of 349 known lipids from 16 classes (see Table 1 for listing of lipid classes). The lipid subclasses in the ADNI serum lipidomics data set used in this study include acylcarnitine, fatty acid, cholesteryl ester, lysophosphatidylcholine, lysophosphatidylethanolamine, phosphatidylcholine, phosphatidylethanolamine, phosphatidylinositol, plasmalogen phosphatidylcholine, plasmalogen phosphatidylethanolamine, ceramide, glucosylceramide, sphingomyelin, diacylglycerol, and triacylglycerol (see Table 1 for listing of lipid classes).

**Table 1:**
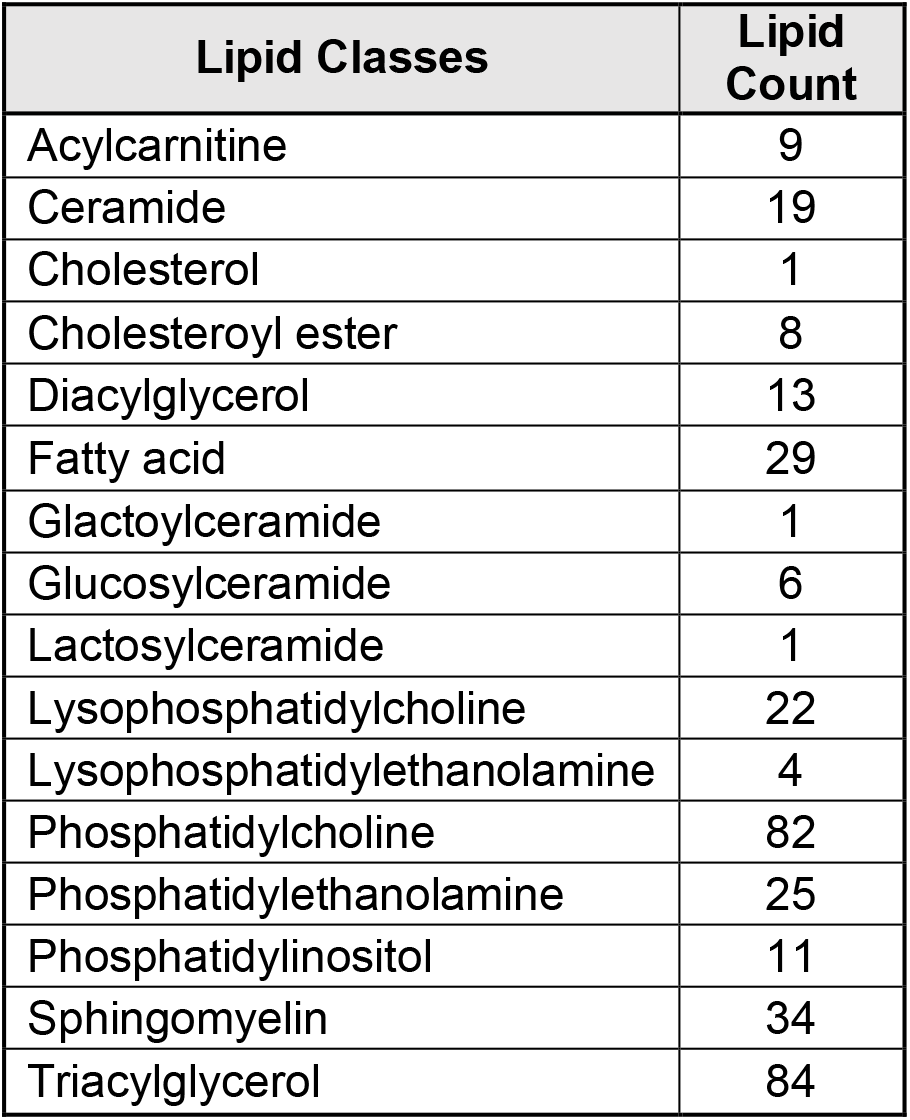
Listing of lipid classes in the current study

| **Lipid Classes** | **Lipid Count** |
| --- | --- |
| Acylcarnitine | 9 |
| Ceramide | 19 |
| Cholesterol | 1 |
| Cholesteroyl ester | 8 |
| Diacylglycerol | 13 |
| Fatty acid | 29 |
| Glactoylceramide | 1 |
| Glucosylceramide | 6 |
| Lactosylceramide | 1 |
| Lysophosphatidylcholine | 22 |
| Lysophosphatidylethanolamine | 4 |
| Phosphatidylcholine | 82 |
| Phosphatidylethanolamine | 25 |
| Phosphatidylinositol | 11 |
| Sphingomyelin | 34 |
| Triacylglycerol | 84 |

### Hearing loss assessment

Hearing was not systematically measured in the ADNI database. Similar to a previous report (Xu et al., 2019), we used subjective hearing loss complaints found in the following datasheets: ADSXLIST.csv, BLSCHECK.csv, INITHEALTH.csv, MEDHIST.csv, NEUROEXM.csv, PHYSICAL.csv, RECBLLOG.csv, RECMHIST.csv. We used the search terms “hear”, “auditory”, “ear”, “deaf”, “presbycusis”, and “HOH (hard of hearing)” and eliminated those reports that were clearly not related to aging-related hearing loss (e.g., skin cancer on ear, earwax, etc.) and eliminated duplicates. Subjects having a hearing complaint are labeled in this study as “hearing loss” or HL. Other subjects are listed as “non-hearing loss” or NHL, notwithstanding the fact that hearing was not objectively measured (see below).

### Statistical Methods

The effect of each individual lipid species on hearing loss in AD subjects was assessed via analysis of covariance (ANCOVA) after adjusting for gender and age as covariates, and log transforming the lipid expression values. Samples with absolute value of studentized residuals from this model exceeding 3 were identified as outliers and excluded from further analysis. The summary measures reported from this analysis include the area under the Receiver Operating Characteristic curve (ROC AUC), covariates-adjusted significance (p-value), and false discovery rate (Benjamini & Hochberg, 1995).

The effect of each of the 16 known lipid classes and 28 empirically derived lipid sets (Barupal et al., 2019) on hearing loss in AD subjects was assessed via “lipid set analysis” (LSA). See Supplementary Table 1 for a listing of the lipids in each of the 28 sets. This LSA analysis of the lipid classes and lipid sets was based on the maxmean statistic of the gene-set-analysis algorithm (Efron & Tibshirani, 2007). which was applied on the residuals from the above ANCOVA model on the individual lipid species to adjust for the effects of age and gender. Individual subject-level standardized composite scores were determined for each lipid class and each lipid set from this algorithm. These scores were then used to assess the effect of each of the lipid classes and lipid sets on hearing loss in AD subjects. The results were summarized in terms of ROC-AUC, covariates-adjusted significance (p-value) and false discovery rate (q-value). Lipid sets with q-value < 0.05 were considered as statistically significant. The corresponding lipid classes and individual lipid specifies with Bonferroni adjusted p-value < 0.05 were highlighted and studied further in terms of their potential connections to hearing loss in AD subjects.

## Results

### Demographics

Data were obtained from 185 subjects with AD. Of the 185, 40 (21.6%) reported hearing loss (HL). HL subjects were not significantly different in age than NHL subjects (HL: 77.2 ± 5.8 years [SD], NHL: 74.8 ± 7.7 years [SD], p > 0.05). HL subjects were more likely to be men than control subjects (NHL = 47% men, HL = 68% men, p < 0.05, Chi-Square). HL and NHL subjects did not differ significantly in average ADAS13 scores (HL: 30.4 ± 8.0 [SD], NHL: 28.8 ± 7.6 [SD], p > 0.05) or body mass index (HL: 26.0 ± 4.1 kg/m^2^ [SD], NHL: 25.3 ± 3.8 kg/m^2^ [SD], p > 0.05, see Table 2).

**Table 2:** Demographic variables. *p<0.05

|  |  | NHL | HL |
| --- | --- | --- | --- |
| **n (# of AD subjects)** |  | 145 | 40<sup>274</sup> |
| **Gender\* (n)** | **F** | 77 | 13 |
|  | **M** | 68 | 27 |
| **Age in years (Mean +/- SD)** |  | 74.8<br>(7.7) | 77.2<br>(5.8) |
| **BMI in kg/m<sup>2</sup> (Mean +/- SD)** |  | 25.32<br>(3.8) | 26<br>(4.1) |
| **ADAS13 (Mean +/- SD)** |  | 28.6<br>(7.6) | 30.4<br>(8) |

### Lipidomic biomarker sets that separate HL from NHL subjects

Levels of 349 lipids were measured across 16 classes. Because levels of many of the lipids are strongly correlated due to co-regulation, and because of the high potential for false discovery when comparing the levels of all 349 lipids, we attempted to reduce the data by grouping the lipids. A previous report measured correlations between all of the serum lipid biomarkers, and using a dynamic clustering algorithm known as dynamicTreeCut, determined that 28 co-regulated sets of lipids were present (Barupal et al., 2019). They also found that many of these lipid sets were associated with either AD diagnosis or AD biomarkers. Although most of the sets were homogeneous (or near-homogeneous) clusters of single lipid types, others comprised a mixture of lipids (see supplementary Table 1 for listing of lipids in each class).

Given the robust performance of these clusters to signal changes in AD biomarkers, we asked whether these same clusters were also associated with the presence of HL. The p- and q- values for the 28 groups of lipids are shown in Table 3. We found that two sets of lipids correlated with the presence of hearing loss: set 23 and set 4, both with p- and q-values below 0.05, with set 23 producing the best performance. We therefore focused on the lipids found in these two sets for subsequent analyses of lipid class and individual lipids.

**Table 3:** Table of lipid sets derived from from Barupal et al. 2019 and their performance in distinguishing HL from NHL subjects. ROC AUC = receiver-operator characteristic area under the curve.

| Lipid Set | Median (HN) | Median (HL) | ROC AUC | p-value (unadj.) | q-value (FDR-BH) |
| --- | --- | --- | --- | --- | --- |
| Set.23 | 0.54 | -2.1 | 0.66 | 0.0006 | 0.0175 |
| Set.4 | 0.16 | -1.06 | 0.64 | 0.0032 | 0.0447 |
| Set.6 | -0.05 | 0.35 | 0.62 | 0.0148 | 0.1332 |
| Set.25 | 1 | -1.3 | 0.62 | 0.019 | 0.1332 |
| Set.3 | 0.37 | -0.74 | 0.6 | 0.0297 | 0.1664 |
| Set.16 | -0.16 | 0.27 | 0.59 | 0.0529 | 0.2171 |
| Set.10 | 0.26 | -1.18 | 0.6 | 0.0543 | 0.2171 |
| Set.14 | 0.51 | -1.16 | 0.6 | 0.0705 | 0.2467 |
| Set.11 | 0.55 | -1.21 | 0.61 | 0.0802 | 0.2495 |
| Set.27 | 0.6 | -1.61 | 0.58 | 0.1025 | 0.2871 |
| Set.19 | 0.2 | -0.35 | 0.58 | 0.1238 | 0.3014 |
| Set.7 | 0.06 | -0.74 | 0.58 | 0.1292 | 0.3014 |
| Set.8 | 0.31 | -0.26 | 0.57 | 0.1694 | 0.3467 |
| Set.28 | 0.4 | -0.12 | 0.56 | 0.1788 | 0.3467 |
| Set.15 | 0.71 | -0.31 | 0.56 | 0.1857 | 0.3467 |
| Set.20 | 0.22 | -0.45 | 0.56 | 0.2013 | 0.3523 |
| Set.13 | -0.67 | -1.3 | 0.56 | 0.2554 | 0.4206 |
| Set.17 | 0.34 | 0.2 | 0.47 | 0.2836 | 0.4387 |
| Set.24 | -0.4 | -1.43 | 0.56 | 0.3135 | 0.4387 |
| Set.2 | -0.2 | 0.25 | 0.47 | 0.3153 | 0.4387 |
| Set.9 | 0.54 | -0.33 | 0.55 | 0.329 | 0.4387 |
| Set.1 | -0.3 | -1.07 | 0.55 | 0.3879 | 0.4937 |
| Set.26 | 0.02 | 0.58 | 0.48 | 0.4248 | 0.5171 |
| Set.22 | -0.18 | -0.35 | 0.53 | 0.4606 | 0.5359 |
| Set.18 | -0.1 | -0.68 | 0.54 | 0.4785 | 0.5359 |
| Set.5 | 0.54 | -0.55 | 0.53 | 0.6265 | 0.6747 |
| Set.21 | -0.21 | 0.32 | 0.5 | 0.8784 | 0.9109 |
| Set.12 | 0.09 | 0.07 | 0.51 | 0.9827 | 0.9827 |

### Lipid classes and individual lipids that separate HL from NHL subjects

Using the biomarker sets to narrow our hypotheses about which lipids exhibit signal changes in hearing, we attempted to determine which lipid classes were most significantly associated with HL. Within the two top sets identified above, 25 lipids in 7 classes were identified, with only the phosphatidylcholine class surviving correction for multiple comparisons (uncorrected p-value = 0.0057, Bonferroni corrected to 0.04). See Figure 1 for boxplots of the 7 biomarker classes comparing HL and NHL subjects See Table 4 for a listing of lipid classes found in sets 4 and 23 and their associated capacity to separate HL from NHL subjects.

**Figure 1:**
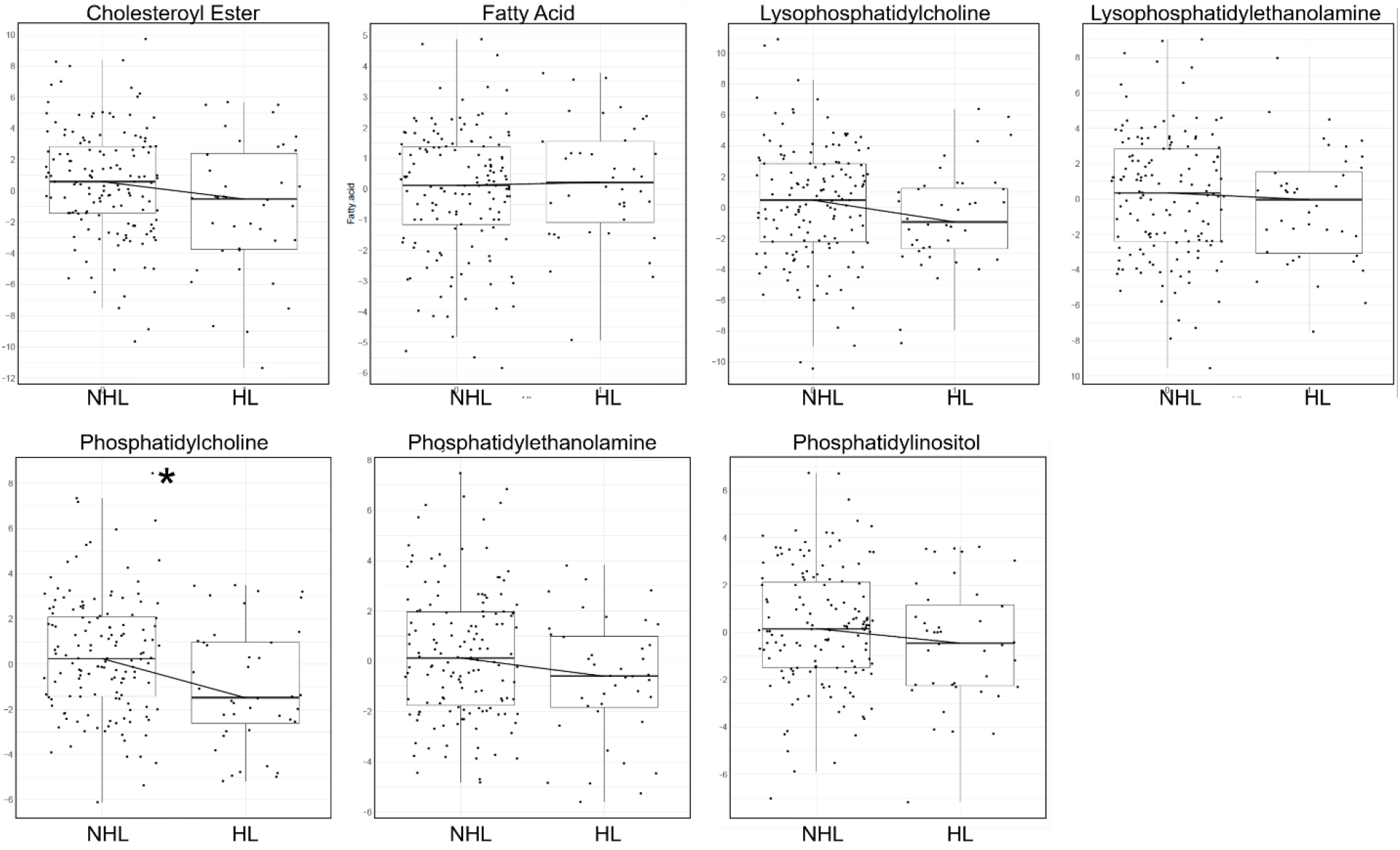
Box plots showing the mean, first and third quartiles of the distributions, demonstrating differences in levels of lipids in the seven classes of lipids identified as parts of sets 4 and 23 from (Barupal et al., 2019), distinguishing between HL and NHL subjects. Shown are standardized values (centered by mean and divided by standard deviation), after adjusting for age and gender as covariates. * Bonferroni-corrected p-value of < 0.05.

**Table 4.** Table of lipid classes derived from sets 4 and 23 from Barupal et al. 2019 and their performance in distinguishing HL from NHL subjects.

| Lipid Class | Median<br>(NHL) | Median<br>(HL) | ROC<br>AUC | p-value |
| --- | --- | --- | --- | --- |
| Phosphatidylcholine | 0.25 | -1.48 | 0.63 | 0.0057 |
| Phosphatidylethanolamine | 0.13 | -0.6 | 0.59 | 0.0216 |
| Cholesteroyl.ester | 0.58 | -0.53 | 0.62 | 0.0239 |
| Phosphatidylinositol | 0.16 | -0.45 | 0.58 | 0.1142 |
| Lysophosphatidylcholine | 0.48 | -0.93 | 0.58 | 0.1255 |
| Lysophosphatidylethanolamine | 0.35 | -0.02 | 0.55 | 0.3185 |
| Fatty.acid | 0.12 | 0.22 | 0.54 | 0.3344 |

Using the 25 lipids identified above, the most commonly-appearing lipid class was phosphatidylcholine (14/25 lipids or 56%), which is significantly greater than the proportion of all tested lipids that were in the phosphatidylcholine class (82/349 lipids or 23.4%, p<0.05, Chi-Squared test). See Table 5 for a listing of individual lipids in sets 4 and 23 and their associated capacity to separate HL from NHL subjects. Both of these analyses point to phosphatidylcholine levels as the main factor distinguishing between HL and NHL subjects.

**Table 5:** Table of lipids derived from sets 4 and 23 from Barupal et al. 2019 and their performance in distinguishing HL from NHL subjects.

| Lipid ID | Lipid Class | Lipid Set | Median (NHL) | Median (HL) | Fold Change (HL/HN) | p-value |
| --- | --- | --- | --- | --- | --- | --- |
| UCD.Lipid.162 | Phosphatidylcholine | Set-23 | 63671 | 52397 | 0.82 | 0.0003 |
| UCD.Lipid.163 | Phosphatidylcholine | Set-23 | 29367 | 24419.5 | 0.83 | 0.0006 |
| UCD.Lipid.148 | Phosphatidylcholine | Set-4 | 5056746 | 4380450.5 | 0.87 | 0.0010 |
| UCD.Lipid.161 | Phosphatidylcholine | Set-23 | 46893 | 39148.5 | 0.83 | 0.0014 |
| UCD.Lipid.164 | Phosphatidylcholine | Set-23 | 20525 | 16734.5 | 0.82 | 0.0033 |
| UCD.Lipid.17 | Cholesteryl ester | Set-4 | 254617 | 198944.5 | 0.78 | 0.0055 |
| UCD.Lipid.150 | Phosphatidylcholine | Set-4 | 58706 | 49712 | 0.85 | 0.0069 |
| UCD.Lipid.406 | Phosphatidylcholine | Set-4 | 130587 | 116420 | 0.89 | 0.0079 |
| UCD.Lipid.128 | Lysophosphatidylcholine | Set-4 | 45190 | 37318.5 | 0.83 | 0.0130 |
| UCD.Lipid.451 | Phosphatidylethanolamine | Set-23 | 5726.33 | 4968 | 0.87 | 0.0149 |
| UCD.Lipid.143 | Phosphatidylcholine | Set-4 | 23780 | 19794 | 0.83 | 0.0150 |
| UCD.Lipid.409 | Phosphatidylcholine | Set-4 | 69527 | 57894 | 0.83 | 0.0163 |
| UCD.Lipid.149 | Phosphatidylcholine | Set-4 | 35516.5 | 27777 | 0.78 | 0.0166 |
| UCD.Lipid.462 | Phosphatidylinositol | Set-4 | 9408 | 8256 | 0.88 | 0.0183 |
| UCD.Lipid.447 | Phosphatidylethanolamine | Set-23 | 12761 | 11301.5 | 0.89 | 0.0197 |
| UCD.Lipid.450 | Phosphatidylethanolamine | Set-23 | 12513.79 | 10888.5 | 0.87 | 0.0217 |
| UCD.Lipid.410 | Phosphatidylcholine | Set-4 | 7910 | 7552 | 0.95 | 0.0310 |
| UCD.Lipid.145 | Phosphatidylcholine | Set-4 | 22891 | 19311 | 0.84 | 0.0329 |
| UCD.Lipid.126 | Lysophosphatidylcholine | Set-4 | 14172.5 | 12585 | 0.89 | 0.0428 |
| UCD.Lipid.399 | Phosphatidylcholine | Set-4 | 98283 | 79011 | 0.80 | 0.0661 |
| UCD.Lipid.16 | Cholesteryl ester | Set-4 | 179126 | 125757 | 0.70 | 0.0846 |
| UCD.Lipid.442 | Phosphatidylethanolamine | Set-4 | 1817.5 | 1492 | 0.82 | 0.1016 |
| UCD.Lipid.381 | Lysophosphatidylethanolamine | Set-4 | 5577.5 | 5206 | 0.93 | 0.2087 |
| UCD.Lipid.517 | Fatty acid | Set-4 | 111966 | 100323 | 0.90 | 0.5032 |
| UCD.Lipid.513 | Fatty acid | Set-4 | 22307 | 21665 | 0.97 | 0.7021 |

### Analysis of non-AD subjects and APOE

Similar analyses were done in subjects with mild cognitive impairment (MCI) and control subjects without memory loss. None of the lipid sets were found to differentiate HL from NHL subjects in either control or MCI cohorts. Subjects across all groups (control, MCI and AD) were also separated based on genotype (having at least one copy of APOE4 or none), and no association was found between genotype and likelihood of HL.

## Discussion

In the current study, 349 serum biomarkers were measured in 185 subjects with AD. Using previously-identified co-regulated sets of biomarkers (Barupal et al., 2019), we found two sets of lipids that were strongly associated with the presence of HL. Within these sets, the most common class of lipids was phosphatidylcholine, and as a class and as individual biomarkers, phosphatidylcholines were found to be significantly diminished in individuals with HL. Similar analyses in non-AD subjects (control and MCI) did not reveal significant associations between lipidomic biomarkers and HL

### Weaknesses in the study

Hearing loss in this study was assessed in a non-systematic way – via reports obtained from the subjects. Previous data have established concordance values between subjective and objective hearing loss ranging from 65-77% depending on demographic factors (Kamil, Genther, & Lin, 2015). Although there are several publicly-available databases that have measured hearing loss objectively (e.g., Baltimore Longitudinal Study of Aging or National Health and Nutrition Examination Survey), these did not systematically measure an extensive panel of lipid biomarkers. Conversely, despite the richness of biomarker data available in ADNI, hearing was not systematically measured. Thus, additional future work in subjects with objectively-measured hearing loss will be required to confirm the associations reported here.

In addition, as an observational study, the current work cannot be used to support the idea that supplementation of phosphatidylcholine can protect against ARHL in subjects with AD. It is possible that phosphatidylcholine levels and ARHL are related by a third, unmeasured, factor. Only a prospective, randomized and blinded trial can determine whether phosphatidylcholine can improve ARHL.

### Phosphatidylcholine, Alzheimer Disease and hearing loss

Phosphatidylcholine is one of the major phospholipids and a fundamental constituent of cell membranes and may activate enzymatic antioxidants situated in the cell membrane. There has also been evidence for disrupted phosphatidylcholine metabolism in AD. For example, the enzymes that break down phosphatidylcholine (phospholipase D and phospholipase A2) are altered with AD (Sanchez-Mejia & Mucke, 2010; Whiley et al., 2014). In addition, low plasma levels phosphatidylcholine docosahexaenoic acid have been associated with the development of AD (Schaefer et al., 2006) as well as thinning of the prefrontal cortex (Zamroziewicz, Zwilling, & Barbey, 2016). With respect to ARHL, phosphatidylcholine’s protective role in hearing loss was suggested by work from Seidman et al, who observed that lecithin (a polyunsaturated phosphatidylcholine) can protect against aging-related hearing loss in rats (Seidman, Khan, Tang, & Quirk, 2002). The antioxidants activated by phosphatidylcholine may protect the cell membrane from damage by reactive oxygen species (Kurutas, 2015) that arise during aging-related cochlear hypoperfusion, which can lead to cochlear degeneration (Gonzalez-Gonzalez, 2017; Seidman, Khan, Dolan, & Quirk, 1996). These data all suggest that phosphatidylcholine levels may be depleted in AD and ARHL.

The lipids measured in this study were extracted from blood samples which brings about the question of the origins of these lipids. Dietary fats are absorbed into the portal system to the liver. In the liver, fatty acids are incorporated into lipoprotein particles which are then released into the bloodstream. Additionally, adipocytes can release stored fatty acids into the blood as lipid levels in the blood decrease. Evidence also suggests that some fatty acids can be synthesized in the brain, but that essential fatty acids still have to be transported across the blood-brain barrier (Bruce, Zsombok, & Eckel, 2017). Additional studies done on adult rats to study the rate of polyunsaturated fatty acid incorporation from plasma into the brain further suggests that this is a dynamic process with active daily turnover (Rapoport, Chang, & Spector, 2001). The exact mechanism behind how fats enter the brain is still unclear. One study performed on cholesterol homeostasis and hearing loss indicates that since the blood-brain barrier prevents the uptake of this lipoprotein from circulation, brain cholesterol is synthesized in astrocytes; further, excess cholesterol is metabolized into 24 (S)- hydroxycholesterol before secretion from the blood-brain barrier to the liver (Malgrange, Varela-Nieto, de Medina, & Paillasse, 2015). Thus, measured lipids in this study likely are likely derived from a variety of sources.

### Conclusion

Thus, in the current study, we observed that in the context of AD, lower serum levels of phosphatidylcholine were associated with ARHL. The fact that this association was found in AD subjects, but not in non-AD subjects, suggests that there is an interaction between the presence of AD and the relationship between phosphatidylcholine and ARHL. Given that AD is associated with diminished brain mitochondrial function and increased levels of lipid peroxidation, it is possible that individuals with AD may not have the metabolic reserve to withstand additional metabolic stressors, such as declining levels of antioxidant molecules such as phosphatidylcholine. These data also suggest that normalizing phosphatidylcholine levels in AD subjects, but not in non-AD subjects, may have a role in for the treatment or prevention of ARHL. Future studies will need to be done to investigate the potential therapeutic role of phosphatidylcholine in this context.

## Acknowledgment

The authors thank Dr. Dinesh Barupal for useful discussions regarding the ADNI lipidomics dataset and Dr. Aditi Das for useful comments on this manuscript.

---

**Supplemental Table 1:** Listing of lipids in all empirically-derived sets from Barupal et al. 2019

| Lipid Set | Lipid Class | Lipid ID | Network Label | FullName | AcylChainInformation | Adduct | RT | MZ |
| --- | --- | --- | --- | --- | --- | --- | --- | --- |
| Set -1 | Triacylglycerol | UCD.Lipid.173 | TG 42:0 | TG (42:0) | TG (14:0_14:0_14:0) | [M+N H4]<sup>+</sup> | 9.4<br>3 | 740.6<br>763 |
| Set -1 | Triacylglycerol | UCD.Lipid.174 | TG 40:0 | TG (40:0) | TG(12:0_12:0_16:0) | [M+N H4]<sup>+</sup> | 9.1 | 712.6<br>419 |
| Set -1 | Triacylglycerol | UCD.Lipid.177 | TG 42:2 | TG (42:2) | TG(12:0_12:0_18:2) | [M+N H4]<sup>+</sup> | 8.4<br>67 | 736.6<br>446 |
| Set -1 | Triacylglycerol | UCD.Lipid.179 | TG 44:0 | TG (44:0) | TG(14:0_14:0_16:0) | [M+N H4]<sup>+</sup> | 10.<br>13 | 768.7<br>037 |
| Set -1 | Triacylglycerol | UCD.Lipid.180 | TG 44:1 | TG (44:1) | TG(12:0_14:0_18:1) | [M+N H4]<sup>+</sup> | 9.4<br>55 | 766.6<br>919 |
| Set -1 | Triacylglycerol | UCD.Lipid.181 | TG 44:2 | TG (44:2) |  | [M+Na]<sup>+</sup> | 9.0<br>33 | 769.6<br>316 |
| Set -1 | Triacylglycerol | UCD.Lipid.182 | TG 46:0 | TG (46:0) | TG(14:0_16:0_16:0) | [M+N H4]<sup>+</sup> | 10.<br>38 | 796.7<br>389 |
| Set -1 | Triacylglycerol | UCD.Lipid.183 | TG 46:1 | TG (46:1) | TG(12:0_16:0_18:1) | [M+N H4]<sup>+</sup> | 9.9<br>44 | 794.7<br>232 |
| Set -1 | Triacylglycerol | UCD.Lipid.184 | TG 46:2 | TG (46:2) | TG(12:0_16:1_18:1) | [M+Na]<sup>+</sup> | 9.5<br>12 | 797.6<br>63 |
| Set -1 | Triacylglycerol | UCD.Lipid.185 | TG 46:3 | TG (46:3) | TG(12:0_16:1_18:2) | [M+Na]<sup>+</sup> | 9.0<br>65 | 795.6<br>473 |
| Set -1 | Triacylglycerol | UCD.Lipid.186 | TG 46:4 A | TG (46:4) A |  | [M+Na]<sup>+</sup> | 8.8 | 793.6<br>322 |
| Set -1 | Triacylglycerol | UCD.Lipid.187 | TG 48:0 | TG (48:0) | TG(16:0_16:0_16:0) | [M+N H4]<sup>+</sup> | 10.<br>81 | 824.7<br>702 |
| Set -1 | Triacylglycerol | UCD.Lipid.188 | TG 48:1 | TG (48:1) | TG(14:0_16:0_18:1) | [M+N H4]<sup>+</sup> | 10.<br>4 | 822.7<br>545 |
| Set -1 | Triacylglycerol | UCD.Lipid.189 | TG 48:2 | TG (48:2) | TG(14:0_16:0_18:2) | [M+N H4]<sup>+</sup> | 9.9<br>93 | 820.7<br>389 |
| Set -1 | Triacylglycerol | UCD.Lipid.192 | TG 49:0 | TG (49:0) | TG(16:0_16:0_17:0) | [M+N H4]<sup>+</sup> | 10.<br>94 | 838.7<br>858 |
| Set -1 | Triacylglycerol | UCD.Lipid.193 | TG 49:1 | TG (49:1) | TG(15:0_16:0_18:1) | [M+N H4]<sup>+</sup> | 10.<br>6 | 836.7<br>702 |
| Set -1 | Triacylglycerol | UCD.Lipid.194 | TG 49:2 | TG (49:2) | TG(15:0_16:0_18:2) | [M+N H4]<sup>+</sup> | 10.<br>18 | 834.7<br>545 |
| Set -1 | Triacylglycerol | UCD.Lipid.196 | TG 50:0 | TG (50:0) | TG(16:0_16:0_18:0) | [M+N H4]<sup>+</sup> | 11.<br>21 | 852.8<br>015 |
| Set -1 | Triacylglycerol | UCD.Lipid.197 | TG 50:1 | TG (50:1) | TG(16:0_16:0_18:1) | [M+N H4]<sup>+</sup> | 10.<br>82 | 850.7<br>858 |
| Set -1 | Triacylglycerol | UCD.Lipid.198 | TG 50:2 | TG (50:2) | TG(16:0_16:1_18:1) | [M+N H4]<sup>+</sup> | 10.<br>42 | 848.7<br>702 |
| Set -1 | Triacylglycerol | UCD.Lipid.202 | TG 50:6 | TG (50:6) | TG(14:0_16:1_20:5) | [M+Na]<sup>+</sup> | 9.1<br>8 | 845.6<br>599 |
| Set -1 | Triacylglycerol | UCD.Lipid.203 | TG 51:1 | TG (51:1) | TG(16:0_17:0_18:1) | [M+N H4]<sup>+</sup> | 10.<br>96 | 864.8<br>015 |
| Set -1 | Triacylglycerol | UCD.Lipid.208 | TG 52:0 | TG (52:0) | TG(16:0_18:0_18:0) | [M+N H4]<sup>+</sup> | 11.<br>58 | 880.8<br>328 |
| Set -1 | Triacylglycerol | UCD.Lipid.209 | TG 52:1 | TG (52:1) | TG(16:0_18:0_18:1) | [M+N H4]<sup>+</sup> | 11.<br>22 | 878.8<br>171 |
| Set -1 | Triacylglycerol | UCD.Lipid.215 | TG 53:1 | TG (53:1) | TG(17:0_18:0_18:1) | [M+N H4]<sup>+</sup> | 11.<br>35 | 892.8<br>328 |
| Set -1 | Triacylglycerol | UCD.Lipid.220 | TG 54:0 | TG (54:0) | TG(18:0_18:0_18:0) | [M+N H4]<sup>+</sup> | 11.<br>92 | 908.8<br>576 |
| Set -1 | Triacylglycerol | UCD.Lipid.221 | TG 54:1 | TG (54:1) | TG(18:0_18:0_18:1) | [M+N H4]<sup>+</sup> | 11.<br>58 | 906.8<br>484 |
| Set -1 | Triacylglycerol | UCD.Lipid.229 | TG 56:2 | TG (56:2) | TG(18:1_18:1_20:0) | [M+N H4]<sup>+</sup> | 11.<br>58 | 932.8<br>641 |
| Set -2 | Cholesteryl ester | UCD.Lipid.10 | CE 16:1 | CE (16:1) | 16:1 Cholesteryl ester | [M+Na]<sup>+</sup> | 10.<br>33 | 645.5<br>581 |
| Set -2 | Lysophosphatidylcholine | UCD.Lipid.119 | LPC 14:0 | LPC (14:0) | LysoPC 14:0 | [M+H]<sup>+</sup> | 0.9<br>81 | 468.3<br>089 |
| Set -2 | Phosphatidylcholine | UCD.Lipid.133 | PC 28:0 | PC (28:0) |  | [M+H]<sup>+</sup> | 4.2<br>39 | 678.5<br>065 |
| Set -2 | Phosphatidylcholine | UCD.Lipi d.134 | PC 30:0 | PC (30:0) | PC(14:0_16:0) | [M+H]<br>+ | 4.8<br>11 | 706.5<br>378 |
| Set -2 | Phosphatidylcholine | UCD.Lipi d.135 | PC 30:1 | PC (30:1) |  | [M+H]<br>+ | 4.3<br>22 | 704.5<br>225 |
| Set -2 | Phosphatidylcholine | UCD.Lipi d.136 | PC 31:0 | PC (31:0) |  | [M+H]<br>+ | 5.1<br>1 | 720.5<br>538 |
| Set -2 | Phosphatidylcholine | UCD.Lipi d.137 | PC 31:1 | PC (31:1) |  | [M+H]<br>+ | 4.8<br>2 | 718.5<br>345 |
| Set -2 | Phosphatidylcholine | UCD.Lipi d.138 | PC 32:3 | PC (32:3) |  | [M+H]<br>+ | 4.8<br>11 | 728.5<br>222 |
| Set -2 | Phosphatidylcholine | UCD.Lipi d.139 | PC 33:0 | PC (33:0) |  | [M+H]<br>+ | 5.6<br>4 | 748.5<br>851 |
| Set -2 | Phosphatidylcholine | UCD.Lipi d.383 | PC 32:1 | PC (32:1) | PC(16:0_16:1) | [M+Ac<br>-H]- | 5.0<br>33 | 790.5<br>6 |
| Set -2 | Phosphatidylcholine | UCD.Lipi d.384 | PC 32:2 | PC (32:2) | PC(14:0_18:2) | [M+Ac<br>-H]- | 4.5<br>67 | 788.5<br>4 |
| Set -2 | Phosphatidylcholine | UCD.Lipi d.385 | PC 33:1 | PC (33:1) | PC(15:0_18:1) | [M+Ac<br>-H]- | 5.3<br>4 | 804.5<br>8 |
| Set -2 | Phosphatidylcholine | UCD.Lipi d.388 | PC 34:1 | PC (34:1) | PC(16:0_18:1) | [M+Ac<br>-H]- | 5.6<br>48 | 818.5<br>9 |
| Set -2 | Phosphatidylcholine | UCD.Lipi d.390 | PC 34:3 | PC (34:3) | PC(16:0_18:3) | [M+Ac<br>-H]- | 4.7<br>5 | 814.5<br>6 |
| Set -2 | Phosphatidylcholine | UCD.Lipi d.391 | PC 34:4 | PC (34:4) | PC(14:0_20:4) | [M+Ac<br>-H]- | 4.4<br>92 | 812.5<br>4 |
| Set -2 | Phosphatidylcholine | UCD.Lipi d.392 | PC 35:1 | PC (35:1) | PC(17:0_18:1) | [M+Ac<br>-H]- | 5.9<br>73 | 832.6<br>1 |
| Set -2 | Phosphatidylcholine | UCD.Lipi d.395 | PC 36:1 | PC (36:1) | PC(18:0_18:1) | [M+Ac<br>-H]- | 6.3<br>14 | 846.6<br>2 |
| Set -2 | Phosphatidylethanolamine | UCD.Lipi d.431 | PE 36:1 | PE (36:1) | PE(18:0_18:1) | [M-<br>H]- | 5.6<br>48 | 744.5<br>5 |
| Set -2 | Phosphatidylethanolamine | UCD.Lipi d.437 | PE 40:6 | PE (40:6) | PE(18:0_22:6) | [M-<br>H]- | 4.9<br>08 | 790.5<br>4 |
| Set -2 | Phosphatidylinositol | UCD.Lipi d.452 | PI 32:1 | PI (32:1) | PI(16:0_16:1) | [M-<br>H]- | 4.0<br>25 | 807.5 |
| Set -2 | Phosphatidylinositol | UCD.Lipi d.458 | PI 36:4 | PI (36:4) | PI(16:0_20:4) | [M-<br>H]- | 4.0<br>92 | 857.5<br>2 |
| Set -3 | Sphingomyelin | UCD.Lipi d.169 | SM d30:1 | SM (d30:1) | SM (d18:1_12:0) | [M+H]<br>+ | 3.6<br>34 | 647.5<br>123 |
| Set -3 | Ceramide | UCD.Lipi d.257 | Cer d32:1 | Ceramide (d32:1) | Cer[NS](d16:1_16:0) | [M+Ac<br>-H]- | 5.0<br>91 | 568.4<br>9 |
| Set -3 | Ceramide | UCD.Lipi d.258 | Cer d33:1 | Ceramide (d33:1) | Cer[NS](d17:1_16:0) | [M+Cl]<br>- | 5.4<br>15 | 558.4<br>7 |
| Set -3 | Ceramide | UCD.Lipi d.260 | Cer d34:1 | Ceramide (d34:1) | Cer[NS](d18:1_16:0) | [M+Ac<br>-H]- | 5.7<br>48 | 596.5<br>3 |
| Set -3 | Ceramide | UCD.Lipi d.261 | Cer d34:2 | Ceramide (d34:2) |  | [M+Ac<br>-H]- | 5.1<br>82 | 594.5<br>1 |
| Set -3 | Ceramide | UCD.Lipi d.262 | Cer d36:1 | Ceramide (d36:1) | Cer[NS](d18:1_18:0) | [M+Cl]<br>- | 6.4<br>55 | 600.5<br>1 |
| Set -3 | Ceramide | UCD.Lipi d.272 | Cer d43:1 | Ceramide (d43:1) | Cer[NS](d19:1_24:0) | [M+Cl]<br>- | 8.6<br>82 | 698.6<br>2 |
| Set -3 | Sphingomyelin | UCD.Lipi d.463 | SM d32:0 | SM (d32:0) |  | [M+Ac<br>-H]- | 4.4<br>67 | 735.5<br>6 |
| Set -3 | Sphingomyelin | UCD.Lipi d.464 | SM d32:1 | SM (d32:1) | SM(d18:1_14:0) | [M+Ac<br>-H]- | 4.2<br>67 | 733.5<br>5 |
| Set -3 | Sphingomyelin | UCD.Lipi d.465 | SM d32:2 | SM (d32:2) |  | [M+Ac<br>-H]- | 3.8<br>18 | 731.5<br>3 |
| Set -3 | Sphingomyelin | UCD.Lipi d.466 | SM d33:1 | SM (d33:1) | SM(d17:1_16:0) | [M+Ac<br>-H]- | 4.5<br>58 | 747.5<br>7 |
| Set -3 | Sphingomyelin | UCD.Lipi d.469 | SM d36:0 | SM (d36:0) |  | [M+Ac<br>-H]- | 5.7<br>65 | 791.6<br>3 |
| Set -3 | Sphingomyelin | UCD.Lipi d.470 | SM d36:1 | SM (d36:1) | SM(d18:1_18:0) | [M+Ac<br>-H]- | 5.5<br>23 | 789.6<br>1 |
| Set -3 | Sphingomyelin | UCD.Lipi d.473 | SM d37:1 | SM (d37:1) |  | [M+Ac<br>-H]- | 5.8<br>73 | 803.6<br>3 |
| Set -3 | Sphingomyelin | UCD.Lipi d.474 | SM d38:0 | SM (d38:0) | SM(d18:0_20:0) | [M+Ac<br>-H]- | 6.4<br>55 | 819.6<br>6 |
| Set -3 | Sphingomyelin | UCD.Lipi d.475 | SM d38:1 | SM (d38:1) | SM(d18:1_20:0) | [M+Ac<br>-H]- | 6.2<br>14 | 817.6<br>4 |
| Set -3 | Sphingomyelin | UCD.Lipi d.477 | SM d39:1 | SM (d39:1) | SM(d16:1_23:0) | [M+Ac-H]- | 6.5<br>55 | 831.6<br>6 |
| Set -3 | Sphingomyelin | UCD.Lipi d.490 | SM d43:1 | SM (d43:1) |  | [M+Ac-H]- | 7.7<br>7 | 887.7<br>2 |
| Set -3 | Sphingomyelin | UCD.Lipi d.491 | SM d43:2 | SM (d43:2) |  | [M+Ac-H]- | 7.0<br>79 | 885.7<br>1 |
| Set -4 | Lysophosphatidylcholine | UCD.Lipi d.126 | LPC 20:5 | LPC (20:5) | LysoPC 20:5 | [M+H]<br>+ | 0.9<br>64 | 542.3<br>239 |
| Set -4 | Lysophosphatidylcholine | UCD.Lipi d.128 | LPC 22:6 | LPC (22:6) | LysoPC 22:6 | [M+H]<br>+ | 1.1<br>55 | 568.3<br>404 |
| Set -4 | Phosphatidylcholine | UCD.Lipi d.143 | PC 36:6 | PC (36:6) |  | [M+H]<br>+ | 4.2<br>31 | 778.5<br>378 |
| Set -4 | Phosphatidylcholine | UCD.Lipi d.145 | PC 37:6 | PC (37:6) |  | [M+H]<br>+ | 4.4<br>96 | 792.5<br>538 |
| Set -4 | Phosphatidylcholine | UCD.Lipi d.148 | PC 38:6 A | PC (38:6) A | PC(16:0_22:6) | [M+H]<br>+ | 4.7<br>78 | 806.5<br>691 |
| Set -4 | Phosphatidylcholine | UCD.Lipi d.149 | PC 38:7 | PC (38:7) |  | [M+H]<br>+ | 4.3<br>14 | 804.5<br>538 |
| Set -4 | Phosphatidylcholine | UCD.Lipi d.150 | PC 39:6 | PC (39:6) |  | [M+H]<br>+ | 5.0<br>27 | 820.5<br>851 |
| Set -4 | Cholesteryl ester | UCD.Lipi d.16 | CE 20:5 | CE (20:5) | 20:5 Cholesteryl ester | [M+Na]<br>+ | 9.7<br>11 | 693.5<br>581 |
| Set -4 | Cholesteryl ester | UCD.Lipi d.17 | CE 22:6 | CE (22:6) | 22:6 Cholesteryl ester | [M+Na]<br>+ | 9.8<br>94 | 719.5<br>738 |
| Set -4 | Lysophosphatidylethanolamine | UCD.Lipi d.381 | LPE 22:6 | LPE (22:6) | LysoPC 22:6 | [M-H]<br>- | 1.2<br>63 | 524.2<br>8 |
| Set -4 | Phosphatidylcholine | UCD.Lipi d.399 | PC 36:5 B | PC (36:5) B | PC(16:1_20:4) | [M+Ac-H]- | 4.6<br>83 | 838.5<br>6 |
| Set -4 | Phosphatidylcholine | UCD.Lipi d.406 | PC 38:6 | PC (38:6) | PC(18:1_20:5)_PC(18:2_20:4) | [M+Ac-H]- | 4.9<br>08 | 864.5<br>8 |
| Set -4 | Phosphatidylcholine | UCD.Lipi d.409 | PC 40:6 B | PC (40:6) B | PC(18:0_22:6) | [M+Ac-H]- | 5.5<br>32 | 892.6<br>1 |
| Set -4 | Phosphatidylcholine | UCD.Lipi d.410 | PC 40:7 | PC (40:7) | PC(18:1_22:6) | [M+Ac-H]- | 4.9<br>83 | 890.5<br>9 |
| Set -4 | Phosphatidylethanolamine | UCD.Lipi d.442 | PE p-36:5 | PE (p-36:5) or PE (o-36:6) | PE(P-16:0_20:5) | [M-H]<br>- | 5.0<br>99 | 720.5 |
| Set -4 | Phosphatidylinositol | UCD.Lipi d.462 | PI 40:6 | PI (40:6) | PI(18:0_22:6) | [M-H]<br>- | 4.5<br>08 | 909.5<br>5 |
| Set -4 | Fatty acid | UCD.Lipi d.513 | FA 20:5 | FA (20:5) | FA (20:5) | [M-H]<br>- | 1.5<br>3 | 301.2<br>2 |
| Set -4 | Fatty acid | UCD.Lipi d.517 | FA 22:6 | FA (22:6) | FA (22:6) | [M-H]<br>- | 1.7<br>2 | 327.2<br>3 |
| Set -5 | Diacylglycerol | UCD.Lipi d.104 | DG 32:1 | DG (32:1) | DG(16:0_16:1) | [M+Na]<br>+ | 6.1<br>43 | 589.4<br>812 |
| Set -5 | Diacylglycerol | UCD.Lipi d.105 | DG 34:1 | DG (34:1) | DG(16:0_18:1) | [M+Na]<br>+ | 6.8<br>92 | 617.5<br>116 |
| Set -5 | Diacylglycerol | UCD.Lipi d.106 | DG 34:2 | DG (34:2) | DG(16:0_18:2) | [M+Na]<br>+ | 6.2<br>7 | 615.4<br>965 |
| Set -5 | Diacylglycerol | UCD.Lipi d.107 | DG 34:3 | DG (34:3) | DG(16:1_18:2) | [M+N H4]<br>+ | 5.6<br>86 | 608.5<br>246 |
| Set -5 | Diacylglycerol | UCD.Lipi d.109 | DG 36:2 | DG (36:2) | DG(18:1_18:1) | [M+N H4]<br>+ | 6.9<br>5 | 638.5<br>718 |
| Set -5 | Diacylglycerol | UCD.Lipi d.110 | DG 36:3 | DG (36:3) | DG(18:1_18:2) | [M+N H4]<br>+ | 6.4<br>2 | 636.5<br>562 |
| Set -5 | Triacylglycerol | UCD.Lipi d.190 | TG 48:3 | TG (48:3) | TG(12:0_18:1_18:2) | [M+Na]<br>+ | 9.5<br>46 | 823.6<br>789 |
| Set -5 | Triacylglycerol | UCD.Lipi d.191 | TG 48:4 | TG (48:4) | TG(12:0_18:2_18:2) | [M+Na]<br>+ | 9.3<br>4 | 821.6<br>597 |
| Set -5 | Triacylglycerol | UCD.Lipi d.195 | TG 49:3 | TG (49:3) | TG(15:0_16:1_18:2) | [M+N H4]<br>+ | 9.7<br>94 | 832.7<br>389 |
| Set -5 | Triacylglycerol | UCD.Lipi d.199 | TG 50:3 | TG (50:3) | TG(16:1_16:1_18:1) | [M+N H4]<br>+ | 10.<br>02 | 846.7<br>545 |
| Set -5 | Triacylglycerol | UCD.Lipi d.200 | TG 50:4 | TG (50:4) | TG(16:1_16:1_18:2) | [M+Na]<br>+ | 9.6<br>29 | 849.6<br>943 |
| Set -5 | Triacylglycerol | UCD.Lipi d.201 | TG 50:5 | TG (50:5) | TG(14:0_18:2_18:3) | [M+Na]<br>+ | 9.2<br>4 | 847.6<br>786 |
| Set -5 | Triacylglycerol | UCD.Lipi d.204 | TG 51:2 | TG (51:2) | TG(16:0_17:0_18:2) | [M+N H4]<br>+ | 10.<br>62 | 862.7<br>858 |
| Set -5 | Triacylglycerol | UCD.Lipi d.205 | TG 51:3 | TG (51:3) | TG(15:0_18:1_18:2) | [M+N H4]+ | 10.25 | 860.7702 |
| Set -5 | Triacylglycerol | UCD.Lipi d.210 | TG 52:2 | TG (52:2) | TG(16:0_18:1_18:1) | [M+N H4]+ | 10.83 | 876.8015 |
| Set -5 | Triacylglycerol | UCD.Lipi d.211 | TG 52:3 | TG (52:3) | TG(16:0_18:1_18:2) | [M+N H4]+ | 10.47 | 874.7858 |
| Set -5 | Triacylglycerol | UCD.Lipi d.216 | TG 53:2 | TG (53:2) | TG(17:0_18:1_18:1) | [M+N H4]+ | 11.01 | 890.8171 |
| Set -5 | Triacylglycerol | UCD.Lipi d.217 | TG 53:3 | TG (53:3) | TG(17:0_18:1_18:2) | [M+N H4]+ | 10.65 | 888.8015 |
| Set -6 | Diacylglycerol | UCD.Lipi d.108 | DG 36:1 | DG (36:1) | DG(18:0_18:1) | [M+N H4]+ | 7.47 | 640.588 |
| Set -6 | Diacylglycerol | UCD.Lipi d.113 | DG 38:0 | DG (38:0) |  | [M+N H4]+ | 7.94 | 670.6086 |
| Set -6 | Diacylglycerol | UCD.Lipi d.114 | DG 38:3 | DG (38:3) | DG(20:1_18:2) | [M+Na ]+ | 6.974 | 669.5653 |
| Set -6 | Diacylglycerol | UCD.Lipi d.116 | DG 38:6 | DG (38:6) | DG(18:2_20:4) | [M+Na ]+ | 5.732 | 663.4959 |
| Set -6 | Lysophosphatidylcholine | UCD.Lipi d.127 | LPC 22:4 | LPC (22:4) | LysoPC 22:4 | [M+H] + | 1.661 | 572.3711 |
| Set -6 | Lysophosphatidylcholine | UCD.Lipi d.129 | LPC o-16:0 | LPC (o-16:0) | LPC (o-16:0) | [M+H] + | 1.752 | 482.3606 |
| Set -6 | Lysophosphatidylcholine | UCD.Lipi d.130 | LPC p-16:0 | LPC (p-16:0) or LPC (o-16:1) | LPC (o-16:1) | [M+H] + | 1.694 | 480.3448 |
| Set -6 | Lysophosphatidylcholine | UCD.Lipi d.131 | LPC p-18:0 | LPC (p-18:0) or LPC (o-18:1) | LPC (o-18:1) | [M+H] + | 1.909 | 508.3762 |
| Set -6 | Triacylglycerol | UCD.Lipi d.175 | TG 40:1 | TG (40:1) |  | [M+N H4]+ | 8.384 | 710.6298 |
| Set -6 | Triacylglycerol | UCD.Lipi d.178 | TG 42:3 | TG (42:3) |  | [M+N H4]+ | 8.24 | 734.6265 |
| Set -6 | Ceramide | UCD.Lipi d.19 | Cer d42:2 | Ceramide (d42:2) | Cer[NS](d18:2_24:0) | [M+H] + | 7.615 | 648.6289 |
| Set -6 | Triacylglycerol | UCD.Lipi d.253 | TG 60:6 | TG (60:6) |  | [M+Na ]+ | 11.27 | 985.8105 |
| Set -6 | Ceramide | UCD.Lipi d.259 | Cer d34:0 | Ceramide (d34:0) | Cer[NDS](d18:0_16:0) | [M+Ac -H]- | 5.998 | 598.54 |
| Set -6 | Phosphatidylethanolamine | UCD.Lipi d.439 | PE p-36:1 | PE (p-36:1) or PE (o-36:2) | PE(P-18:0_18:1) | [M-H]- | 6.838 | 728.56 |
| Set -6 | Fatty acid | UCD.Lipi d.511 | FA 20:3 | FA (20:3) | FA (20:3) | [M-H]- | 2.5 | 305.25 |
| Set -6 | Fatty acid | UCD.Lipi d.512 | FA 20:4 | FA (20:4) | FA (20:4) | [M-H]- | 1.93 | 303.23 |
| Set -6 | Fatty acid | UCD.Lipi d.515 | FA 22:1 | FA (22:1) | FA (22:1) | [M-H]- | 3.9 | 337.31 |
| Set -6 | Acylcarnitine | UCD.Lipi d.9 | AC C18:3 | Acylcarnitine C18:3 | Acylcarnitine 18:3 | [M+H] + | 1.13 | 422.3265 |
| Set -7 | Phosphatidylcholine | UCD.Lipi d.142 | PC 36:4 A | PC (36:4) A | PC(18:2_18:2) | [M+H] + | 4.637 | 782.5694 |
| Set -7 | Glucosylceramide | UCD.Lipi d.368 | GlcCer d24:1-2OH | GlcCer(d14:1(4E)/20:0(2OH)) |  | [M+Cl] - | 4.966 | 750.53 |
| Set -7 | Phosphatidylcholine | UCD.Lipi d.382 | PC 32:0 | PC (32:0) | PC(16:0_16:0) | [M+Ac -H]- | 5.573 | 792.58 |
| Set -7 | Phosphatidylcholine | UCD.Lipi d.386 | PC 33:2 | PC (33:2) | PC(16:0_18:2) | [M+Ac -H]- | 4.858 | 802.56 |
| Set -7 | Phosphatidylcholine | UCD.Lipi d.387 | PC 34:0 | PC (34:0) | PC(16:0_18:0) | [M+Ac -H]- | 6.239 | 820.61 |
| Set -7 | Phosphatidylcholine | UCD.Lipi d.389 | PC 34:2 | PC (34:2) | PC(16:0_18:2) | [M+Ac -H]- | 5.166 | 816.58 |
| Set -7 | Phosphatidylcholine | UCD.Lipi d.393 | PC 35:2 | PC (35:2) | PC(17:0_18:2) | [M+Ac -H]- | 5.48 | 830.59 |
| Set -7 | Phosphatidylcholine | UCD.Lipi d.396 | PC 36:2 | PC (36:2) | PC(18:0_18:2) | [M+Ac -H]- | 5.806 | 844.61 |
| Set -7 | Phosphatidylcholine | UCD.Lipi d.400 | PC 37:2 | PC (37:2) | PC(19:0_18:2) | [M+Ac -H]- | 6.14 | 858.62 |
| Set -7 | Phosphatidylethanolamine | UCD.Lipi d.434 | PE 38:2 | PE (38:2) | PE(20:0_18:2) | [M-H]- | 5.806 | 770.57 |
| Set -7 | Phosphatidylethanolamine | UCD.Lipid.443 | PE p-38:2 | PE (p-38:2) or PE (o-38:3) |  | [M-H]<sup>-</sup> | 6.98 | 754.58 |
| Set -7 | Phosphatidylethanolamine | UCD.Lipid.448 | PE p-40:4 | PE (p-40:4) or PE (o-40:5) | PE(P-20:0_20:4) | [M-H]<sup>-</sup> | 6.87 | 778.58 |
| Set -7 | Phosphatidylinositol | UCD.Lipid.453 | PI 34:1 | PI (34:1) | PI(16:0_18:1) | [M-H]<sup>-</sup> | 4.483 | 835.53 |
| Set -7 | Phosphatidylinositol | UCD.Lipid.454 | PI 34:2 | PI (34:2) | PI(16:0_18:2) | [M-H]<sup>-</sup> | 4.158 | 833.52 |
| Set -7 | Phosphatidylinositol | UCD.Lipid.455 | PI 36:1 | PI (36:1) | PI(18:0_18:1) | [M-H]<sup>-</sup> | 5.198 | 863.57 |
| Set -7 | Phosphatidylinositol | UCD.Lipid.456 | PI 36:2 | PI (36:2) | PI(18:0_18:2) | [M-H]<sup>-</sup> | 4.732 | 861.55 |
| Set -7 | Phosphatidylinositol | UCD.Lipid.457 | PI 36:3 | PI (36:3) | PI(16:0_20:3) | [M-H]<sup>-</sup> | 4.225 | 859.53 |
| Set -8 | Lysophosphatidylcholine | UCD.Lipid.120 | LPC 15:0 | LPC (15:0) | LysoPC 15:0 | [M+H]<sup>+</sup> | 1.206 | 482.3241 |
| Set -8 | Lysophosphatidylcholine | UCD.Lipid.121 | LPC 17:1 | LPC (17:1) | LysoPC 17:1 | [M+H]<sup>+</sup> | 1.345 | 508.3401 |
| Set -8 | Lysophosphatidylcholine | UCD.Lipid.122 | LPC 18:0 | LPC (18:0) | LysoPC 18:0 | [M+H]<sup>+</sup> | 2.258 | 524.3708 |
| Set -8 | Lysophosphatidylcholine | UCD.Lipid.123 | LPC 18:3 | LPC (18:3) | LysoPC 18:3 | [M+H]<sup>+</sup> | 0.981 | 518.3241 |
| Set -8 | Lysophosphatidylcholine | UCD.Lipid.124 | LPC 20:0 | LPC (20:0) | LysoPC 20:0 | [M+H]<sup>+</sup> | 3.062 | 552.4021 |
| Set -8 | Lysophosphatidylcholine | UCD.Lipid.369 | LPC 16:0 | LPC (16:0) | LysoPC 16:0 | [M+Ac-H]<sup>-</sup> | 1.563 | 554.35 |
| Set -8 | Lysophosphatidylcholine | UCD.Lipid.370 | LPC 16:1 | LPC (16:1) | LysoPC 16:1 | [M+Ac-H]<sup>-</sup> | 1.18 | 552.33 |
| Set -8 | Lysophosphatidylcholine | UCD.Lipid.371 | LPC 18:0 A | LPC (18:0) A | LysoPC 18:0 | [M+Ac-H]<sup>-</sup> | 2.145 | 582.38 |
| Set -8 | Lysophosphatidylcholine | UCD.Lipid.372 | LPC 18:1 | LPC (18:1) | LysoPC 18:1 | [M+Ac-H]<sup>-</sup> | 1.713 | 580.36 |
| Set -8 | Lysophosphatidylcholine | UCD.Lipid.373 | LPC 18:2 | LPC (18:2) | LysoPC 18:2 | [M+Ac-H]<sup>-</sup> | 1.313 | 578.35 |
| Set -8 | Lysophosphatidylcholine | UCD.Lipid.374 | LPC 20:1 | LPC (20:1) | LysoPC 20:1 | [M+Ac-H]<sup>-</sup> | 2.47 | 608.39 |
| Set -8 | Lysophosphatidylcholine | UCD.Lipid.375 | LPC 20:2 | LPC (20:2) | LysoPC 20:2 | [M+Ac-H]<sup>-</sup> | 1.896 | 606.38 |
| Set -8 | Lysophosphatidylcholine | UCD.Lipid.376 | LPC 20:3 | LPC (20:3) | LysoPC 20:3 | [M+Ac-H]<sup>-</sup> | 1.505 | 604.36 |
| Set -8 | Lysophosphatidylcholine | UCD.Lipid.377 | LPC 22:5 | LPC (22:5) | LysoPC 22:5 | [M+Ac-H]<sup>-</sup> | 1.388 | 628.36 |
| Set -8 | Lysophosphatidylethanolamine | UCD.Lipid.378 | LPE 16:0 | LPE (16:0) | LysoPE 16:0 | [M-H]<sup>-</sup> | 1.596 | 452.28 |
| Set -8 | Lysophosphatidylethanolamine | UCD.Lipid.379 | LPE 18:2 | LPE (18:2) | LysoPE 18:2 | [M-H]<sup>-</sup> | 1.347 | 476.28 |
| Set -9 | Phosphatidylcholine | UCD.Lipid.132 | PC 27:0-CHO | PC (16:0/9:0(CHO)) | PC (16:0_9:0(CHO)) | [M+H]<sup>+</sup> | 2.374 | 650.4391 |
| Set -9 | Triacylglycerol | UCD.Lipid.176 | TG 42:1 | TG (42:1) | TG(12:0_12:0_18:1) | [M+Na]<sup>+</sup> | 8.967 | 743.6162 |
| Set -9 | Fatty acid | UCD.Lipid.493 | FA 11:0 | FA (11:0) | FA (11:0) | [M-H]<sup>-</sup> | 0.82 | 185.15 |
| Set -9 | Fatty acid | UCD.Lipid.494 | FA 12:0 | FA (12:0) | FA (12:0) | [M-H]<sup>-</sup> | 1.06 | 199.17 |
| Set -9 | Fatty acid | UCD.Lipid.495 | FA 13:0 | FA (13:0) | FA (13:0) | [M-H]<sup>-</sup> | 1.3 | 213.19 |
| Set -9 | Fatty acid | UCD.Lipid.496 | FA 14:0 | FA (14:0) | FA (14:0) | [M-H]<sup>-</sup> | 1.63 | 227.2 |
| Set -9 | Fatty acid | UCD.Lipid.498 | FA 15:0 | FA (15:0) | FA (15:0) | [M-H]<sup>-</sup> | 2.02 | 241.22 |
| Set -9 | Fatty acid | UCD.Lipid.499 | FA 15:1 | FA (15:1) | FA (15:1) | [M-H]<sup>-</sup> | 1.5 | 239.2 |
| Set -9 | Fatty acid | UCD.Lipid.500 | FA 16:0 | FA (16:0) | FA (16:0) | [M-H]<sup>-</sup> | 2.47 | 255.23 |
| Set -9 | Fatty acid | UCD.Lipid.502 | FA 17:0 | FA (17:0) | FA (17:0) | [M-H]<sup>-</sup> | 2.96 | 269.25 |
| Set -9 | Fatty acid | UCD.Lipid.504 | FA 18:0 | FA (18:0) | FA (18:0) | [M-H]<sup>-</sup> | 3.26 | 283.26 |
| Set -9 | Fatty acid | UCD.Lipi d.508 | FA 20:0 | FA (20:0) | FA (20:0) | [M-H]<sup>-</sup> | 3.8<br>5 | 311.3 |
| Set -9 | Fatty acid | UCD.Lipi d.514 | FA 22:0 | FA (22:0) | FA (22:0) | [M-H]<sup>-</sup> | 4.5 | 339.3<br>3 |
| Set -9 | Fatty acid | UCD.Lipi d.518 | FA 24:0 | FA (24:0) | FA (24:0) | [M-H]<sup>-</sup> | 5.2<br>3 | 367.3<br>6 |
| Set -9 | Fatty acid | UCD.Lipi d.520 | FA 26:0 | FA (26:0) | FA (26:0) | [M-H]<sup>-</sup> | 6.0<br>2 | 395.3<br>9 |
| Set -9 | Fatty acid | UCD.Lipi d.521 | FA 28:0 | FA (28:0) | FA (28:0) | [M-H]<sup>-</sup> | 6.8<br>2 | 423.4<br>2 |
| Set -10 | Sphingomyelin | UCD.Lipi d.171 | SM d40:2 | SM (d40:2) |  | [M+H]<sup>+</sup> | 6.0<br>38 | 785.6<br>531 |
| Set -10 | Phosphatidylinositol | UCD.Lipi d.461 | PI 38:5 | PI (38:5) | PI(18:1_20:4) | [M-H]<sup>-</sup> | 4.1<br>42 | 883.5<br>3 |
| Set -10 | Sphingomyelin | UCD.Lipi d.468 | SM d34:2 | SM (d34:2) | SM(d18:2_16:0) | [M+Ac-H]<sup>-</sup> | 4.3<br>59 | 759.5<br>7 |
| Set -10 | Sphingomyelin | UCD.Lipi d.471 | SM d36:2 | SM (d36:2) |  | [M+Ac-H]<sup>-</sup> | 4.9<br>74 | 787.6 |
| Set -10 | Sphingomyelin | UCD.Lipi d.472 | SM d36:3 | SM (d36:3) |  | [M+Ac-H]<sup>-</sup> | 4.5<br>08 | 785.5<br>8 |
| Set -10 | Sphingomyelin | UCD.Lipi d.476 | SM d38:2 | SM (d38:2) |  | [M+Ac-H]<sup>-</sup> | 5.6<br>23 | 815.6<br>3 |
| Set -10 | Sphingomyelin | UCD.Lipi d.478 | SM d39:2 | SM (d39:2) |  | [M+Ac-H]<sup>-</sup> | 5.9<br>88 | 829.6<br>4 |
| Set -10 | Sphingomyelin | UCD.Lipi d.479 | SM d40:0 | SM (d40:0) |  | [M+Ac-H]<sup>-</sup> | 7.1<br>38 | 847.6<br>9 |
| Set -10 | Sphingomyelin | UCD.Lipi d.480 | SM d40:1 | SM (d40:1) | SM(d18:1_22:0) | [M+Ac-H]<sup>-</sup> | 6.8<br>88 | 845.6<br>7 |
| Set -10 | Sphingomyelin | UCD.Lipi d.481 | SM d40:2 A | SM (d40:2) A |  | [M+Ac-H]<sup>-</sup> | 6.2<br>05 | 843.6<br>6 |
| Set -10 | Sphingomyelin | UCD.Lipi d.482 | SM d40:2 B | SM (d40:2) B | SM(d18:2_22:0) | [M+Ac-H]<sup>-</sup> | 6.3<br>06 | 843.6<br>6 |
| Set -10 | Sphingomyelin | UCD.Lipi d.483 | SM d40:3 | SM (d40:3) |  | [M+Ac-H]<sup>-</sup> | 5.6<br>4 | 841.6<br>4 |
| Set -10 | Sphingomyelin | UCD.Lipi d.484 | SM d41:1 | SM (d41:1) | SM(d18:1_23:0) | [M+Ac-H]<sup>-</sup> | 7.2<br>29 | 859.6<br>9 |
| Set -10 | Sphingomyelin | UCD.Lipi d.485 | SM d41:2 | SM (d41:2) | SM(d18:2_23:0) | [M+Ac-H]<sup>-</sup> | 6.6<br>5 | 857.6<br>7 |
| Set -10 | Sphingomyelin | UCD.Lipi d.486 | SM d42:0 | SM (d42:0) |  | [M+Ac-H]<sup>-</sup> | 7.7<br>94 | 875.7<br>2 |
| Set -10 | Sphingomyelin | UCD.Lipi d.487 | SM d42:1 | SM (d42:1) | SM(d18:1_24:0) | [M+Ac-H]<sup>-</sup> | 7.5<br>62 | 873.7<br>1 |
| Set -11 | Glactoylceramide | UCD.Lipi d.117 | LacCer d34:1 | Gal-Gal-Cer(d18:1/16:0) or Lactosylceramide(d18:1/16:0) |  | [M+H]<sup>+</sup> | 4.7<br>95 | 862.6<br>25 |
| Set -11 | Lactosylceramide | UCD.Lipi d.118 | LacCer d32:2 | Lactosylceramide (d18:1/24:1(15Z)) |  | [M+H]<sup>+</sup> | 6.6<br>85 | 972.7<br>346 |
| Set -11 | Cholesteryl ester | UCD.Lipi d.12 | CE 18:2 | CE (18:2) | 18:2 Cholesteryl ester | [M+N H4]<sup>+</sup> | 10.<br>4 | 666.6<br>184 |
| Set -11 | Phosphatidylcholine | UCD.Lipi d.155 | PC o-34:0 | PC (o-34:0) |  | [M+H]<sup>+</sup> | 6.4<br>78 | 748.6<br>204 |
| Set -11 | Phosphatidylcholine | UCD.Lipi d.156 | PC p-32:1 | PC (p-32:1) or PC (o-32:2) |  | [M+H]<sup>+</sup> | 4.9<br>1 | 716.5<br>626 |
| Set -11 | Phosphatidylcholine | UCD.Lipi d.157 | PC p-34:1 | PC (p-34:1) or PC (o-34:2) B | PC(P-16:0_18:1) | [M+H]<sup>+</sup> | 5.7<br>9 | 744.5<br>908 |
| Set -11 | Phosphatidylcholine | UCD.Lipi d.160 | PC p-40:1 | PC (p-40:1) or PC (o-40:2) |  | [M+H]<sup>+</sup> | 6.6<br>37 | 828.6<br>842 |
| Set -11 | Sphingomyelin | UCD.Lipi d.170 | SM d34:0 | SM (d34:0) |  | [M+H]<sup>+</sup> | 4.9<br>6 | 705.5<br>905 |
| Set -11 | Glucosylceramide | UCD.Lipi d.363 | GlcCer d38:1 | GlcCer (d38:1) |  | [M+Ac-H]<sup>-</sup> | 6.4<br>89 | 814.6<br>4 |
| Set -11 | Glucosylceramide | UCD.Lipi d.364 | GlcCer d40:1 | GlcCer (d40:1) | GlcCer[NS](d18:1_22:0) | [M+Ac-H]<sup>-</sup> | 7.1<br>63 | 842.6<br>7 |
| Set -11 | Glucosylceramide | UCD.Lipi d.365 | GlcCer d41:1 | GlcCer (d41:1) |  | [M+Ac-H]<sup>-</sup> | 7.4<br>95 | 856.6<br>9 |
| Set -11 | Glucosylceramide | UCD.Lipi d.366 | GlcCer d42:1 | GlcCer (d42:1) | GlcCer[NS](d18:1_24:0) | [M+Ac-H]<sup>-</sup> | 7.8<br>2 | 870.7 |
| Set -11 | Glucosylceramide | UCD.Lipi d.367 | GlcCer d42:2 | GlcCer (d42:2) | GlcCer[NS](d18:1_24:1) | [M+Ac -H]- | 7.1 38 | 868.6 9 |
| Set -11 | Phosphatidylcholine | UCD.Lipi d.412 | PC o-32:0 | PC (o-32:0) |  | [M+Ac -H]- | 5.9 73 | 778.5 9 |
| Set -11 | Phosphatidylcholine | UCD.Lipi d.414 | PC p-34:0 | PC (p-34:0) or PC (o-34:1) | PC(P-18:0_16:0) | [M+Ac -H]- | 6.0 39 | 804.6 1 |
| Set -11 | Sphingomyelin | UCD.Lipi d.467 | SM d34:1 | SM (d34:1) | SM(d18:1_16:0) | [M+Ac -H]- | 4.8 66 | 761.5 8 |
| Set -12 | Diacylglycerol | UCD.Lipi d.111 | DG 36:4 | DG (36:4) | DG(18:2_18:2) | [M+N H4]+ | 5.8 15 | 634.5 404 |
| Set -12 | Diacylglycerol | UCD.Lipi d.112 | DG 36:5 | DG (36:5) | DG(18:2_18:3) | [M+N H4]+ | 5.6 5 | 632.5 248 |
| Set -12 | Triacylglycerol | UCD.Lipi d.206 | TG 51:4 | TG (51:4) | TG(15:0_18:2_18:2) | [M+Na ]+ | 9.8 77 | 863.7 099 |
| Set -12 | Triacylglycerol | UCD.Lipi d.207 | TG 51:5 | TG (51:5) | TG(15:0_18:2_18:3) | [M+N H4]+ | 9.7 7 | 856.7 347 |
| Set -12 | Triacylglycerol | UCD.Lipi d.212 | TG 52:4 | TG (52:4) | TG(16:1_18:1_18:2) | [M+N H4]+ | 10. 1 | 872.7 702 |
| Set -12 | Triacylglycerol | UCD.Lipi d.213 | TG 52:5 | TG (52:5) | TG(16:0_18:2_18:3) | [M+N H4]+ | 9.7 28 | 870.7 545 |
| Set -12 | Triacylglycerol | UCD.Lipi d.214 | TG 52:6 | TG (52:6) | TG(16:1_18:2_18:3) | [M+Na ]+ | 9.3 5 | 873.6 943 |
| Set -12 | Triacylglycerol | UCD.Lipi d.218 | TG 53:4 | TG (53:4) | TG(17:1_18:1_18:2) | [M+N H4]+ | 10. 32 | 886.7 858 |
| Set -12 | Triacylglycerol | UCD.Lipi d.219 | TG 53:5 | TG (53:5) | TG(17:1_18:2_18:2) | [M+N H4]+ | 9.9 02 | 884.7 702 |
| Set -12 | Triacylglycerol | UCD.Lipi d.223 | TG 54:3 | TG (54:3) | TG(18:0_18:1_18:2) | [M+N H4]+ | 10. 86 | 902.8 171 |
| Set -12 | Triacylglycerol | UCD.Lipi d.224 | TG 54:4 | TG (54:4) | TG(18:1_18:1_18:2) | [M+N H4]+ | 10. 49 | 900.8 015 |
| Set -12 | Triacylglycerol | UCD.Lipi d.225 | TG 54:5 | TG (54:5) | TG(18:1_18:2_18:2) | [M+Na ]+ | 10. 12 | 903.7 412 |
| Set -12 | Triacylglycerol | UCD.Lipi d.226 | TG 54:6 | TG (54:6) | TG(18:1_18:2_18:3) | [M+Na ]+ | 9.7 45 | 901.7 256 |
| Set -12 | Triacylglycerol | UCD.Lipi d.227 | TG 54:8 | TG (54:8) | TG(18:2_18:3_18:3) | [M+Na ]+ | 8.9 4 | 897.6 943 |
| Set -13 | Triacylglycerol | UCD.Lipi d.228 | TG 56:1 | TG (56:1) | TG(18:0_18:1_20:0) | [M+N H4]+ | 11. 85 | 934.8 797 |
| Set -13 | Triacylglycerol | UCD.Lipi d.237 | TG 57:1 | TG (57:1) |  | [M+N H4]+ | 12 | 948.8 863 |
| Set -13 | Triacylglycerol | UCD.Lipi d.238 | TG 57:2 | TG (57:2) |  | [M+N H4]+ | 11. 88 | 946.8 718 |
| Set -13 | Triacylglycerol | UCD.Lipi d.239 | TG 58:1 | TG (58:1) | TG(18:0_18:1_22:0) | [M+N H4]+ | 11. 99 | 962.9 11 |
| Set -13 | Triacylglycerol | UCD.Lipi d.241 | TG 58:2 | TG (58:2) | TG(18:1_18:1_22:0) | [M+N H4]+ | 11. 85 | 960.8 954 |
| Set -13 | Triacylglycerol | UCD.Lipi d.242 | TG 58:3 | TG (58:3) | TG(18:1_20:1_20:1) | [M+N H4]+ | 11. 6 | 958.8 797 |
| Set -13 | Triacylglycerol | UCD.Lipi d.247 | TG 59:2 | TG (59:2) |  | [M+N H4]+ | 12 | 974.9 031 |
| Set -13 | Triacylglycerol | UCD.Lipi d.248 | TG 59:3 | TG (59:3) |  | [M+N H4]+ | 11. 89 | 972.8 871 |
| Set -13 | Triacylglycerol | UCD.Lipi d.250 | TG 60:2 | TG (60:2) | TG(18:1_20:1_22:0) | [M+N H4]+ | 11. 99 | 988.9 267 |
| Set -13 | Triacylglycerol | UCD.Lipi d.251 | TG 60:3 | TG (60:3) | TG(18:1_20:1_22:1) | [M+N H4]+ | 11. 95 | 986.9 034 |
| Set -13 | Triacylglycerol | UCD.Lipi d.252 | TG 60:4 | TG (60:4) | TG(18:1_20:2_22:1) | [M+N H4]+ | 11. 78 | 984.8 877 |
| Set -13 | Triacylglycerol | UCD.Lipi d.254 | TG 62:3 | TG (62:3) |  | [M+N H4]+ | 12. 04 | 1014. 934 |
| Set -13 | Triacylglycerol | UCD.Lipi d.255 | TG 62:4 | TG (62:4) |  | [M+N H4]+ | 11. 94 | 1012. 918 |
| Set -13 | Triacylglycerol | UCD.Lipi d.256 | TG 64:4 | TG (64:4) |  | [M+N H4]+ | 12. 04 | 1040. 949 |
| Set -14 | Ceramide | UCD.Lipi d.18 | Cer d41:1 | Ceramide (d18:1/23:0) | Cer[NS](d18:1_23:0) | [M+H] + | 7.9 63 | 636.6 289 |
| Set -14 | Ceramide | UCD.Lipi d.263 | Cer d38:1 | Ceramide (d38:1) | Cer[NS](d18:1_20:0) | [M+Cl] - | 7.1 63 | 628.5 4 |
| Set -14 | Ceramide | UCD.Lipi d.264 | Cer d39:1 | Ceramide (d39:1) | Cer[NS](d18:1_22:0) | [M+Cl]<br>- | 7.5<br>12 | 642.5<br>6 |
| Set -14 | Ceramide | UCD.Lipi d.265 | Cer d40:0 | Ceramide (d40:0) | Cer[NDS](d18:0_22:0) | [M+Ac-H]<br>- | 8.0<br>61 | 682.6<br>3 |
| Set -14 | Ceramide | UCD.Lipi d.266 | Cer d40:1 | Ceramide (d40:1) | Cer[NS](d18:1_22:0) | [M+Cl]<br>- | 7.8<br>36 | 656.5<br>8 |
| Set -14 | Ceramide | UCD.Lipi d.267 | Cer d40:2 | Ceramide (d40:2) | Cer[NS](d18:2_22:0) | [M+Cl]<br>- | 7.1<br>85 | 654.5<br>6 |
| Set -14 | Ceramide | UCD.Lipi d.268 | Cer d41:1 | Ceramide (d41:1) | Cer[NS](d18:1_23:0) | [M+Cl]<br>- | 8.1<br>61 | 670.5<br>9 |
| Set -14 | Ceramide | UCD.Lipi d.269 | Cer d42:0 | Ceramide (d42:0) | Cer[NDS](d18:0_24:0) | [M+Cl]<br>- | 8.6<br>85 | 686.6<br>2 |
| Set -14 | Ceramide | UCD.Lipi d.270 | Cer d42:1 | Ceramide (d42:1) | Cer[NS](d18:1_24:0) | [M+Ac-H]<br>- | 8.4<br>69 | 708.6<br>5 |
| Set -14 | Ceramide | UCD.Lipi d.271 | Cer d42:2 A | Ceramide (d42:2) A | Cer[NS](d18:1_24:1) | [M+Cl]<br>- | NA | NA |
| Set -14 | Ceramide | UCD.Lipi d.273 | Cer d44:1 | Ceramide (d44:1) |  | [M+Cl]<br>- | 9.0<br>68 | 712.6<br>4 |
| Set -15 | Phosphatidylcholine | UCD.Lipi d.159 | PC p-38:4 | PC (p-38:4) or PC (o-38:5) B | PC(P-18:0_20:4) | [M+H]<br>+ | 5.8<br>23 | 794.6<br>054 |
| Set -15 | Phosphatidylethanolamine | UCD.Lipi d.167 | PE 38:4 | PE (38:4) | PE(18:0_20:4) | [M+H]<br>+ | 5.7<br>23 | 768.5<br>56 |
| Set -15 | Phosphatidylcholine | UCD.Lipi d.413 | PC p-32:0 | PC (p-32:0) or PC (o-32:1) | PC(P-16:0_16:0) | [M+Ac-H]<br>- | 5.8<br>81 | 776.5<br>8 |
| Set -15 | Phosphatidylcholine | UCD.Lipi d.415 | PC p-34:1 | PC (p-34:1) or PC (o-34:2) A | PC(P-16:0_18:1) | [M+Ac-H]<br>- | 5.5<br>4 | 802.6 |
| Set -15 | Phosphatidylcholine | UCD.Lipi d.416 | PC p-34:2 | PC (p-34:2) or PC (o-34:3) | PC(P-16:0_18:2) | [M+Ac-H]<br>- | 5.4<br>4 | 800.5<br>8 |
| Set -15 | Phosphatidylcholine | UCD.Lipi d.417 | PC p-36:1 | PC (p-36:1) or PC (o-36:2) | PC(P-18:0_18:1) | [M+Ac-H]<br>- | 6.2 | 830.6<br>3 |
| Set -15 | Phosphatidylcholine | UCD.Lipi d.418 | PC p-36:2 | PC (p-36:2) or PC (o-36:3) | PC(P-18:0_18:2) | [M+Ac-H]<br>- | 5.5<br>9 | 828.6<br>1 |
| Set -15 | Phosphatidylcholine | UCD.Lipi d.419 | PC p-36:3 | PC (p-36:3) or PC (o-36:4) | PC(P-16:0_20:3) | [M+Ac-H]<br>- | 5.4<br>4 | 826.6 |
| Set -15 | Phosphatidylcholine | UCD.Lipi d.420 | PC p-36:4 | PC (p-36:4) or PC (o-36:5) | PC(P-16:0_20:4) | [M+Ac-H]<br>- | 5.3<br>32 | 824.5<br>8 |
| Set -15 | Phosphatidylcholine | UCD.Lipi d.422 | PC p-38:4 | PC (p-38:4) or PC (o-38:5) A | PC(P-18:0_20:4) | [M+Ac-H]<br>- | 5.4<br>82 | 852.6<br>1 |
| Set -15 | Phosphatidylcholine | UCD.Lipi d.423 | PC p-38:5 | PC (p-38:5) or PC (o-38:6) | PC(P-18:0_20:5) | [M+Ac-H]<br>- | 5.2<br>57 | 850.6 |
| Set -16 | Acylcarnitine | UCD.Lipi d.1 | AC C8:0 | Acylcarnitine C8:0 | Acylcarnitine 8:0 | [M+H]<br>+ | 0.5<br>5 | 288.2<br>169 |
| Set -16 | Acylcarnitine | UCD.Lipi d.2 | AC C8:1 | Acylcarnitine C8:1 | Acylcarnitine 8:1 | [M+H]<br>+ | 0.5<br>2 | 286.2<br>013 |
| Set -16 | Cholesterol | UCD.Lipi d.20 | Cholesterol | Cholesterol |  | [M-H2O+H]<br>+ | 4.8<br>61 | 369.3<br>516 |
| Set -16 | Acylcarnitine | UCD.Lipi d.3 | AC C10:1 | Acylcarnitine C10:1 | Acylcarnitine 10:1 | [M+H]<br>+ | 0.5<br>9 | 314.2<br>326 |
| Set -16 | Acylcarnitine | UCD.Lipi d.4 | AC C12:0 | Acylcarnitine C12:0 | Acylcarnitine 12:0 | [M+H]<br>+ | 0.7<br>4 | 344.2<br>801 |
| Set -16 | Acylcarnitine | UCD.Lipi d.5 | AC C16:0 | Acylcarnitine C16:0 | Acylcarnitine 16:0 | [M+H]<br>+ | 1.5<br>86 | 400.3<br>42 |
| Set -16 | Fatty acid | UCD.Lipi d.519 | FA 24:1 | FA (24:1) | FA (24:1) | [M-H]<br>- | 4.5<br>2 | 365.3<br>4 |
| Set -16 | Acylcarnitine | UCD.Lipi d.6 | AC C18:0 | Acylcarnitine C18:0 | Acylcarnitine 18:0 | [M+H]<br>+ | 2.3<br>4 | 428.3<br>727 |
| Set -16 | Acylcarnitine | UCD.Lipi d.7 | AC C18:1 | Acylcarnitine C18:1 | Acylcarnitine 18:1 | [M+H]<br>+ | 1.7<br>43 | 426.3<br>578 |
| Set -16 | Acylcarnitine | UCD.Lipi d.8 | AC C18:2 | Acylcarnitine C18:2 | Acylcarnitine 18:2 | [M+H]<br>+ | 1.3<br>29 | 424.3<br>425 |
| Set -17 | Fatty acid | UCD.Lipi d.497 | FA 14:1 | FA (14:1) | FA (14:1) | [M-H]<br>- | 1.2 | 225.1<br>9 |
| Set -17 | Fatty acid | UCD.Lipi d.501 | FA 16:1 | FA (16:1) | FA (16:1) | [M-H]<br>- | 1.8<br>2 | 253.2<br>2 |
| Set -17 | Fatty acid | UCD.Lipi d.503 | FA 17:1 | FA (17:1) | FA (17:1) | [M-H]<br>- | 2.2<br>2 | 267.2<br>3 |
| Set -17 | Fatty acid | UCD.Lipi d.505 | FA 18:1 | FA (18:1) | FA (18:1) | [M-H]<sup>-</sup> | 2.7 | 281.25 |
| Set -17 | Fatty acid | UCD.Lipi d.506 | FA 18:2 | FA (18:2) | FA (18:2) | [M-H]<sup>-</sup> | 2.05 | 279.23 |
| Set -17 | Fatty acid | UCD.Lipi d.507 | FA 18:3 | FA (18:3) | FA (18:3) | [M-H]<sup>-</sup> | 1.623 | 277.22 |
| Set -17 | Fatty acid | UCD.Lipi d.509 | FA 20:1 | FA (20:1) | FA (20:1) | [M-H]<sup>-</sup> | 3.35 | 309.28 |
| Set -17 | Fatty acid | UCD.Lipi d.510 | FA 20:2 | FA (20:2) | FA (20:2) | [M-H]<sup>-</sup> | 2.91 | 307.26 |
| Set -17 | Fatty acid | UCD.Lipi d.516 | FA 22:2 | FA (22:2) | FA (22:2) | [M-H]<sup>-</sup> | 3.47 | 335.3 |
| Set -18 | Diacylglycerol | UCD.Lipi d.115 | DG 38:5 | DG (38:5) | DG(18:1_20:4) | [M+N H4]<sup>+</sup> | 6.246 | 660.5562 |
| Set -18 | Triacylglycerol | UCD.Lipi d.222 | TG 54:2 | TG (54:2) | TG(18:0_18:1_18:1) | [M+N H4]<sup>+</sup> | 11.22 | 904.8328 |
| Set -18 | Triacylglycerol | UCD.Lipi d.230 | TG 56:3 | TG (56:3) | TG(18:0_18:1_20:2) | [M+N H4]<sup>+</sup> | 11.22 | 930.8484 |
| Set -18 | Triacylglycerol | UCD.Lipi d.231 | TG 56:4 | TG (56:4) | TG(18:0_18:1_20:3) | [M+N H4]<sup>+</sup> | 10.9 | 928.8328 |
| Set -18 | Triacylglycerol | UCD.Lipi d.232 | TG 56:5 | TG (56:5) | TG(18:0_18:1_20:4) | [M+N H4]<sup>+</sup> | 10.78 | 926.8171 |
| Set -18 | Triacylglycerol | UCD.Lipi d.233 | TG 56:6 | TG (56:6) | TG(18:1_18:1_20:4) | [M+N H4]<sup>+</sup> | 10.34 | 924.8015 |
| Set -18 | Triacylglycerol | UCD.Lipi d.234 | TG 56:7 | TG (56:7) | TG(18:1_18:2_20:4) | [M+N H4]<sup>+</sup> | 9.968 | 922.7858 |
| Set -18 | Triacylglycerol | UCD.Lipi d.243 | TG 58:4 | TG (58:4) | TG(18:1_18:2_22:1) | [M+N H4]<sup>+</sup> | 11.25 | 956.8641 |
| Set -18 | Triacylglycerol | UCD.Lipi d.244 | TG 58:6 | TG (58:6) | TG(18:0_18:1_22:5) | [M+N H4]<sup>+</sup> | 10.67 | 952.8328 |
| Set -19 | Phosphatidylcholine | UCD.Lipi d.147 | PC 38:5 B | PC (38:5) B | PC(16:0_22:5) | [M+H]<sup>+</sup> | 5.168 | 808.5848 |
| Set -19 | Phosphatidylcholine | UCD.Lipi d.151 | PC 40:5 B | PC (40:5) B | PC(18:0_22:5) | [M+H]<sup>+</sup> | 5.806 | 836.6164 |
| Set -19 | Phosphatidylcholine | UCD.Lipi d.152 | PC 40:6 A | PC (40:6) A | PC(18:0_22:6) | [M+H]<sup>+</sup> | 5.151 | 834.6007 |
| Set -19 | Phosphatidylcholine | UCD.Lipi d.153 | PC 42:5 | PC (42:5) |  | [M+H]<sup>+</sup> | 6.072 | 864.645 |
| Set -19 | Phosphatidylcholine | UCD.Lipi d.154 | PC 42:6 | PC (42:6) |  | [M+H]<sup>+</sup> | 5.773 | 862.6311 |
| Set -19 | Phosphatidylcholine | UCD.Lipi d.405 | PC 38:5 A | PC (38:5) A | PC(18:1_20:4) | [M+Ac -H]<sup>-</sup> | 5.107 | 866.59 |
| Set -19 | Phosphatidylcholine | UCD.Lipi d.408 | PC 40:5 A | PC (40:5) A | PC(18:0_22:5) | [M+Ac -H]<sup>-</sup> | 5.74 | 894.62 |
| Set -19 | Phosphatidylcholine | UCD.Lipi d.411 | PC 40:8 | PC (40:8) | PC(18:2_22:6) | [M+Ac -H]<sup>-</sup> | 4.533 | 888.58 |
| Set -19 | Phosphatidylinositol | UCD.Lipi d.460 | PI 38:4 | PI (38:4) | PI(18:0_20:4) | [M-H]<sup>-</sup> | 4.606 | 885.55 |
| Set -20 | Phosphatidylcholine | UCD.Lipi d.158 | PC p-38:2 | PC (p-38:2) or PC (o-38:3) |  | [M+H]<sup>+</sup> | 6.212 | 798.6359 |
| Set -20 | Phosphatidylcholine | UCD.Lipi d.165 | PC p-42:3 | PC (p-42:3) or PC (o-42:4) |  | [M+H]<sup>+</sup> | 7.224 | 852.6859 |
| Set -20 | Phosphatidylcholine | UCD.Lipi d.421 | PC p-38:3 | PC (p-38:3) or PC (o-38:4) | PC(P-18:0_20:3) | [M+Ac -H]<sup>-</sup> | 6.098 | 854.63 |
| Set -20 | Phosphatidylcholine | UCD.Lipi d.424 | PC p-40:3 | PC (p-40:3) or PC (o-40:4) |  | [M+Ac -H]<sup>-</sup> | 6.763 | 882.66 |
| Set -20 | Phosphatidylcholine | UCD.Lipi d.425 | PC p-40:4 | PC (p-40:4) or PC (o-40:5) | PC(P-20:0_20:4) | [M+Ac -H]<sup>-</sup> | 6.106 | 880.64 |
| Set -20 | Phosphatidylcholine | UCD.Lipi d.426 | PC p-42:4 | PC (p-42:4) or PC (o-42:5) | PC(P-22:0_20:4) | [M+Ac -H]<sup>-</sup> | 6.747 | 908.67 |
| Set -20 | Phosphatidylcholine | UCD.Lipi d.427 | PC p-42:5 | PC (p-42:5) or PC (o-42:6) |  | [M+Ac -H]<sup>-</sup> | 6.188 | 906.66 |
| Set -20 | Phosphatidylcholine | UCD.Lipi d.428 | PC p-44:4 | PC (p-44:4) or PC (o-44:5) |  | [M+Ac -H]<sup>-</sup> | 7.387 | 936.71 |
| Set -21 | Lysophosphatidylcholine | UCD.Lipi d.125 | LPC 20:4 | LPC (20:4) | LysoPC 20:4 | [M+H]<sup>+</sup> | 1.205 | 544.3403 |
| Set -21 | Phosphatidylcholine | UCD.Lipi d.146 | PC 38:4 B | PC (38:4) B |  | [M+H]<sup>+</sup> | 6.146 | 810.6004 |
| Set -21 | Lysophosphatidylethanolamine | UCD.Lipid.380 | LPE 20:4 | LPE (20:4) | LysoPE 20:4 | [M-H]<sup>-</sup> | 1.3<br>13 | 500.2<br>8 |
| Set -21 | Phosphatidylcholine | UCD.Lipid.394 | PC 35:4 | PC (35:4) | PC(15:0_20:4) | [M+Ac-H]<sup>-</sup> | 4.7<br>65 | 826.5<br>6 |
| Set -21 | Phosphatidylcholine | UCD.Lipid.398 | PC 36:4 B | PC (36:4) B | PC(16:0_20:4) | [M+Ac-H]<sup>-</sup> | 5.0<br>74 | 840.5<br>8 |
| Set -21 | Phosphatidylcholine | UCD.Lipid.401 | PC 37:4 | PC (37:4) | PC(17:0_20:4) | [M+Ac-H]<sup>-</sup> | 5.3<br>9 | 854.5<br>9 |
| Set -21 | Phosphatidylcholine | UCD.Lipid.404 | PC 38:4 A | PC (38:4) A | PC(18:0_20:4) | [M+Ac-H]<sup>-</sup> | 5.7<br>06 | 868.6<br>1 |
| Set -21 | Phosphatidylcholine | UCD.Lipid.407 | PC 40:4 | PC (40:4) | PC(18:0_22:4) | [M+Ac-H]<sup>-</sup> | 6.1<br>72 | 896.6<br>4 |
| Set -22 | Phosphatidylethanolamine | UCD.Lipid.168 | PE p-34:1 | PE (p-34:1) or PE (o-34:2) | PE(16:0_18:1) | [M+H]<sup>+</sup> | 6.1<br>6 | 702.5<br>404 |
| Set -22 | Phosphatidylethanolamine | UCD.Lipid.438 | PE p-34:2 | PE (p-34:2) or PE (o-34:3) | PE(P-16:0_18:2) | [M-H]<sup>-</sup> | 5.6<br>4 | 698.5<br>1 |
| Set -22 | Phosphatidylethanolamine | UCD.Lipid.440 | PE p-36:2 | PE (p-36:2) or PE (o-36:3) | PE(P-18:0_18:2) | [M-H]<sup>-</sup> | 6.3<br>14 | 726.5<br>4 |
| Set -22 | Phosphatidylethanolamine | UCD.Lipid.441 | PE p-36:4 | PE (p-36:4) or PE (o-36:5) | PE(P-16:0_20:4) | [M-H]<sup>-</sup> | 5.5<br>23 | 722.5<br>1 |
| Set -22 | Phosphatidylethanolamine | UCD.Lipid.444 | PE p-38:3 | PE (p-38:3) or PE (o-38:4) |  | [M-H]<sup>-</sup> | 6.4<br>8 | 752.5<br>6 |
| Set -22 | Phosphatidylethanolamine | UCD.Lipid.445 | PE p-38:4 | PE (p-38:4) or PE (o-38:5) | PE(P-18:0_20:4) | [M-H]<sup>-</sup> | 6.1<br>89 | 750.5<br>4 |
| Set -22 | Phosphatidylethanolamine | UCD.Lipid.446 | PE p-38:5 | PE (p-38:5) or PE (o-38:6) | PE(P-18:0_20:5) | [M-H]<sup>-</sup> | 5.5<br>82 | 748.5<br>3 |
| Set -22 | Phosphatidylethanolamine | UCD.Lipid.449 | PE p-40:5 | PE (p-40:5) or PE (o-40:6) | PE(P-18:0_22:5) | [M-H]<sup>-</sup> | 6.2<br>14 | 776.5<br>6 |
| Set -23 | Phosphatidylcholine | UCD.Lipid.161 | PC p-40:5 | PC (p-40:5) or PC (o-40:6) |  | [M+H]<sup>+</sup> | 5.7<br>4 | 820.6<br>215 |
| Set -23 | Phosphatidylcholine | UCD.Lipid.162 | PC p-40:6 | PC (p-40:6) or PC (o-40:7) A |  | [M+H]<sup>+</sup> | 5.1<br>6 | 818.6<br>058 |
| Set -23 | Phosphatidylcholine | UCD.Lipid.163 | PC p-40:6 | PC (p-40:6) or PC (o-40:7) B |  | [M+H]<sup>+</sup> | 5.6<br>25 | 818.6<br>058 |
| Set -23 | Phosphatidylcholine | UCD.Lipid.164 | PC p-40:7 | PC (p-40:7) or PC (o-40:8) |  | [M+H]<sup>+</sup> | 5.0<br>85 | 816.5<br>902 |
| Set -23 | Phosphatidylethanolamine | UCD.Lipid.447 | PE p-38:6 | PE (p-38:6) or PE (o-38:7) | PE(16:0_22:6) | [M-H]<sup>-</sup> | 5.3<br>4 | 746.5<br>1 |
| Set -23 | Phosphatidylethanolamine | UCD.Lipid.450 | PE p-40:6 | PE (p-40:6) or PE (o-40:7) | PE(P-18:0_22:6) | [M-H]<sup>-</sup> | 5.9<br>98 | 774.5<br>4 |
| Set -23 | Phosphatidylethanolamine | UCD.Lipid.451 | PE p-40:7 | PE (p-40:7) or PE (o-40:8) | PE(P-18:1_22:6) | [M-H]<sup>-</sup> | 5.4<br>07 | 772.5<br>3 |
| Set -24 | Phosphatidylethanolamine | UCD.Lipid.166 | PE 36:4 | PE (36:4) | PE(16:0_20:4) | [M+H]<sup>+</sup> | 5.2<br>5 | 740.5<br>184 |
| Set -24 | Phosphatidylcholine | UCD.Lipid.429 | PE 34:1 | PE (34:1) | PE(16:0_18:1) | [M-H]<sup>-</sup> | 5.8<br>31 | 716.5<br>2 |
| Set -24 | Phosphatidylethanolamine | UCD.Lipid.430 | PE 34:2 | PE (34:2) | PE(16:0_18:2) | [M-H]<sup>-</sup> | 5.3<br>32 | 714.5<br>1 |
| Set -24 | Phosphatidylethanolamine | UCD.Lipid.432 | PE 36:2 | PE (36:2) | PE(18:0_18:2) | [M-H]<sup>-</sup> | 5.9<br>89 | 742.5<br>4 |
| Set -24 | Phosphatidylethanolamine | UCD.Lipid.433 | PE 36:3 | PE (36:3) | PE(18:1_18:2) | [M-H]<sup>-</sup> | 5.3<br>99 | 740.5<br>2 |
| Set -24 | Phosphatidylethanolamine | UCD.Lipid.435 | PE 38:4 B | PE (38:4) B | PE(18:0_20:4) | [M-H]<sup>-</sup> | 5.8<br>81 | 766.5<br>4 |
| Set -24 | Phosphatidylethanolamine | UCD.Lipid.436 | PE 38:6 | PE (38:6) | PE(16:0_22:6) | [M-H]<sup>-</sup> | 5.0<br>74 | 762.5<br>1 |
| Set -25 | Phosphatidylcholine | UCD.Lipid.140 | PC 35:3 | PC (35:3) |  | [M+H]<sup>+</sup> | 4.8<br>61 | 770.5<br>694 |
| Set -25 | Phosphatidylcholine | UCD.Lipid.141 | PC 36:3 B | PC (36:3) B | PC(18:1_18:2) | [M+H]<sup>+</sup> | 5.1<br>84 | 784.5<br>848 |
| Set -25 | Phosphatidylcholine | UCD.Lipid.144 | PC 37:3 | PC (37:3) |  | [M+H]<sup>+</sup> | 5.4<br>25 | 798.6<br>004 |
| Set -25 | Phosphatidylcholine | UCD.Lipid.397 | PC 36:3 A | PC (36:3) A | PC(18:1_18:2) | [M+Ac-H]<sup>-</sup> | 5.2<br>3 | 842.5<br>9 |
| Set -25 | Phosphatidylcholine | UCD.Lipid.402 | PC 38:2 | PC (38:2) | PC(20:0_18:2) | [M+Ac-H]<sup>-</sup> | 6.3<br>89 | 872.6<br>4 |
| Set -25 | Phosphatidylcholine | UCD.Lipid.403 | PC 38:3 | PC (38:3) | PC(18:0_20:3) | [M+Ac-H]<sup>-</sup> | 5.9<br>81 | 870.6<br>2 |
| Set -25 | Phosphatidylinositol | UCD.Lipi d.459 | PI 38:3 | PI (38:3) | PI(18:0_20:3) | [M-H]<sup>-</sup> | 4.9 | 887.57 |
| Set -26 | Triacylglycerol | UCD.Lipi d.235 | TG 56:8 | TG (56:8) | TG(18:1_18:2_20:5) | [M+Na]<sup>+</sup> | 9.769 | 925.7256 |
| Set -26 | Triacylglycerol | UCD.Lipi d.236 | TG 56:9 | TG (56:9) | TG(18:2_18:2_20:5) | [M+Na]<sup>+</sup> | 9.48 | 923.7057 |
| Set -26 | Triacylglycerol | UCD.Lipi d.240 | TG 58:10 | TG (58:10) | TG(18:2_18:2_22:6) | [M+Na]<sup>+</sup> | 9.396 | 949.7256 |
| Set -26 | Triacylglycerol | UCD.Lipi d.245 | TG 58:8 | TG (58:8) | TG(18:1_18:2_22:5) | [M+N H4]<sup>+</sup> | 10.17 | 948.8015 |
| Set -26 | Triacylglycerol | UCD.Lipi d.246 | TG 58:9 | TG (58:9) | TG(18:2_18:2_22:5) | [M+Na]<sup>+</sup> | 9.794 | 951.7412 |
| Set -26 | Triacylglycerol | UCD.Lipi d.249 | TG 60:11 | TG (60:11) |  | [M+N H4]<sup>+</sup> | 9.645 | 970.7859 |
| Set -27 | Sphingomyelin | UCD.Lipi d.172 | SM d42:2 | SM (d42:2) | SM(d18:1_24:1) | [M+H]<sup>+</sup> | 6.685 | 813.6873 |
| Set -27 | Sphingomyelin | UCD.Lipi d.488 | SM d42:2 A | SM (d42:2) A | SM(d18:1_24:1) | [M+Ac-H]<sup>-</sup> | 6.863 | 871.69 |
| Set -27 | Sphingomyelin | UCD.Lipi d.489 | SM d42:3 | SM (d42:3) | SM(d18:2_24:1) | [M+Ac-H]<sup>-</sup> | 6.289 | 869.68 |
| Set -27 | Sphingomyelin | UCD.Lipi d.492 | SM d44:2 | SM (d44:2) |  | [M+Ac-H]<sup>-</sup> | 7.528 | 899.72 |
| Set -28 | Cholesteryl ester | UCD.Lipi d.11 | CE 18:1 | CE (18:1) | 18:1 Cholesteryl ester | [M+Na]<sup>+</sup> | 10.88 | 673.5894 |
| Set -28 | Cholesteryl ester | UCD.Lipi d.13 | CE 18:3 | CE (18:3) | 18:3 Cholesteryl ester | [M+Na]<sup>+</sup> | 9.985 | 669.5581 |
| Set -28 | Cholesteryl ester | UCD.Lipi d.14 | CE 20:3 | CE (20:3) | 20:3 Cholesteryl ester | [M+N H4]<sup>+</sup> | 10.49 | 692.634 |
| Set -28 | Cholesteryl ester | UCD.Lipi d.15 | CE 20:4 | CE (20:4) | 20:4 Cholesteryl ester | [M+Na]<sup>+</sup> | 10.16 | 695.5738 |

